# Heterogeneous Viral Strategies Promote Coexistence in Virus-Microbe Systems

**DOI:** 10.1101/297127

**Authors:** Hayriye Gulbudak, Joshua S. Weitz

**Affiliations:** Department of Mathematics, University of Louisiana at Lafayette, Lafayette, Louisiana, USA; School of Biological Sciences, Georgia Institute of Technology, Atlanta, GA, USA; School of Physics, Georgia Institute of Technology, Atlanta, GA, USA

## Abstract

Viruses of microbes, including bacterial viruses (phage), archaeal viruses, and eukaryotic viruses, can influence the fate of individual microbes and entire populations. Here, we model distinct modes of virus-host interactions and study their impact on the abundance and diversity of both viruses and their microbial hosts. We consider two distinct viral populations infecting the same microbial population via two different strategies: lytic and chronic. A lytic strategy corresponds to viruses that exclusively infect and lyse their hosts to release new virions. A chronic strategy corresponds to viruses that infect hosts and then continually release new viruses via a budding process without cell lysis. The chronic virus can also be passed on to daughter cells during cell division. The long-term association of virus and microbe in the chronic mode drives differences in selective pressures with respect to the lytic mode. We utilize invasion analysis of the corresponding nonlinear differential equation model to study the ecology and evolution of heterogenous viral strategies. We first investigate stability of equilibria, and characterize oscillatory and bistable dynamics in some parameter regions. Then, we derive fitness quantities for both virus types and investigate conditions for competitive exclusion and coexistence. In so doing we find unexpected results, including a regime in which the chronic virus requires the lytic virus for survival and invasion.

## I. INTRODUCTION

Viruses shape the population and evolutionary dynamics of microbes [1–7]. For example, microbes can evolve resistance to infection and also acquire immunity to infection via intracellular defense mechanisms including resistance-modification, CRISPR-Cas, BREX and other systems [9, 10]. The abundance and diversity of viruses, along with microbes whom they infect, are well documented. The interactions between viruses and their microbial hosts play a central role in maintenance of virus-microbial biodiversity and variation in functional traits within virus-microbial communities [1, 12, 14].

Viruses can interact with microbial hosts given divergent strategies, including lytic, lysogenic, and chronic modes of infection. Virulent viruses reproduce exclusively by lysing their hosts. In contrast, temperate viruses can lyse their hosts or integrate their genomes with that of their hosts thereby forming a lysogen, i.e., replicating with the same cell lineage [18, 19]. In some cases, a virus can be temperate for one host and virulent for another [20]. Despite there is a growing interest in filamentous phage [17], the chronic infection mode is less well characterized. In a chronic infection, the viral genome neither integrates into the host genome nor induces lysis. Instead, the virus forms a persistent infection where progeny may bud from the cell as the viral genome is passed to daughter cells asymmetrically after division [1].

Differing viral infection modes can also influence host fitness in distinct ways, which may in turn impact the virus-microbe community composition. For example temperate viruses may enable infected cells to resist subsequent infections via a mechanism of ‘super infection immunity’[11]. Yet, lysogenic modes of infection can also have fitness costs for microbial hosts [31]. In contrast, viruses that can only lyse their hosts act as predators of microbes and can decrease overall host population fitness [1]. However, the evolution of resistance to lytic viruses by microbial hosts can incur fitness costs [8]. Chronic infections may also prevent infected cells against subsequent lytic viral infection (via superinfection exclusion). However such modes may also carry costs evidenced by the variety of resistance mechanisms that hosts display against them [1].

Mathematical models have provided some insights into how distinct infection modes affect the dynamics of interacting viral populations and, in turn, shape evolutionary dynamics. In doing so, the bulk of evolutionary dynamic models of virus-microbe interactions focus on the lytic mode. Many studies have addressed how inter- and intraspecies interactions shape the evolution of virulent viruses and their microbial hosts [14, 22, 23]. For example, researchers have studied the effect of limited resources on the population dynamics of microbial or virus strains in the paradigm of coexistence or exclusion of species [24, 25]. Competing bacteria and viruses may persist together as a result of predator-mediated coexistence when multiple bacteria and viral species interact given emergent variation in resistance and virulence [4, 26-28]. A few studies have expanded beyond this lytic paradigm to address the question of why viruses evolve lysogeny in the first place [32]. Recent models of evolution in fluctuating environments suggest that the switch between lysis and lysogeny may be shaped by long-term optimization of fitness, leading to the evolution of bet-hedging like strategies[21]. Recent discoveries of the chronic mode of infection raises a related class of questions: what are the costs and benefits of lytic and chronic modes when viruses utilizing these heterogeneous strategies compete for a common host?

In this paper, we analyze the population and evolutionary dynamics of viruses and microbial hosts given concurrent lytic and chronic modes of infection. To do so, we develop a nonlinear model of interactions between lytic and chronic viruses and a common host population. We first analyze infection dynamics arising from chronic infections, and identify the emergence of complex bifurcations as the derived viral reproduction number varies. The reproduction number and bifurcations are distinct from those in models of lytic infection dynamics. The broader conceptual foundations underlying the viral reproduction number analyzed for a spectrum of viral strategies is presented in our companion work [29]. In this paper, we then focus on the system with lytic and chronically infecting virus strains interacting together. Through analytical and numerical methods, we determine the parameter regimes and underlying mechanisms leading to coexistence or exclusion of the distinct virus types, including a regime in which the chronic virus - counterintuitively - requires the virulent lytic virus for survival and invasion.

## II. METHODS

Here we propose a model of virus-host interactions. The model describes the dynamics of interactions between multiple virus strains and one type host. In the model, *S*(*t*), *I*(*t*), *C*(*t*) represent the number of susceptible, infected, chronically infected cells at time t, respectively, and *V_L_*(*t*), *V_C_*(*t*) depict the density of free lytic and chronically infecting virus particles, respectively. We assume that during infective process, the lytic (or chronically infecting) virus is absorbed and infects the susceptible cells with a rate *ϕ* (or 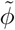). If lytic infection takes place, host cells lyse with a rate η and produce *β viable* virions. If a chronic infection takes place, chronically infecting viruses produce progeny, which are slowly budded off the cell with a rate *α* or passed down to daughter cells and divide with a rate 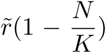. Chronic infections are assumed not to lead to direct cellular lysis as a means to release virus particles. Cells die with a rate d and viruses decay with a rate *μ*. We assume that chronic infection is costly, i.e., such that chronically infected cells die with a rate 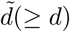.

A nonlinear differential equation system, describing the interactions and the dynamics of host and virus populations, can be written as follows:

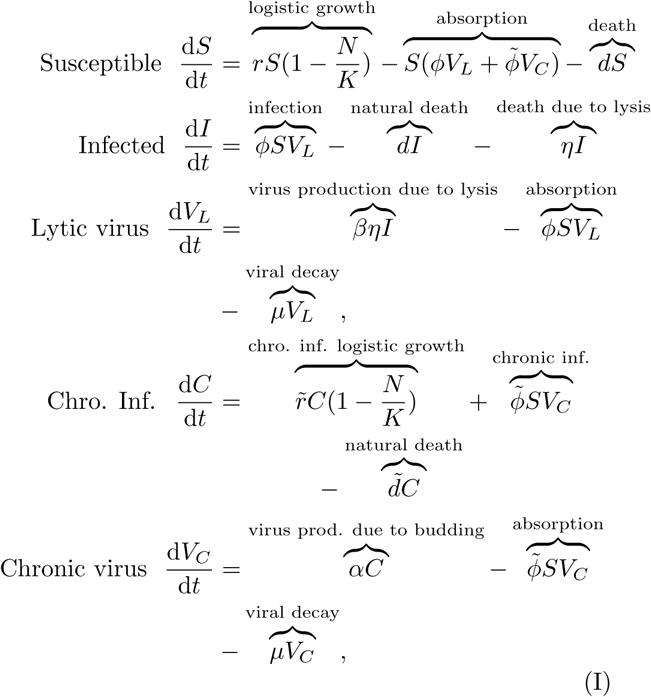

where *N* is the total number of cells, *N* = *S* + *I* + *C*.

We do not consider co-infection or superinfection due to cross immunity provided by each viral infection against the other strain (superinfection exclusion). In particular, we assume that when infection of a host cell with two virus particles takes place, competition between virus particles in a cell for a limited amount of key enzyme more often results in exclusion of all but one of them (*key enzyme* hypothesis [20]). The logistic term in the cell growth rate depicts the competition between infected and susceptible cells for limited resources with carrying capacity *K*.

In this paper, we are interested in studying the outcome of competing lytic and chronic strains on virus-host ecology and evolution in model (I). To do so, we first explore and summarize infection dynamics under each type of virus infection mode separately in next section (section (III)), which will also facilitate analysis of the multi-strain model in Section IV.

## III. INFECTION DYNAMICS

### A. Lytic Infection Dynamics

First we reduce the system of equations (I) to a threedimensional system by taking *V_C_* (*t*) = 0 and *C*(*t*) = 0, in other words, we consider the following subsystem with only lytic viral infection:

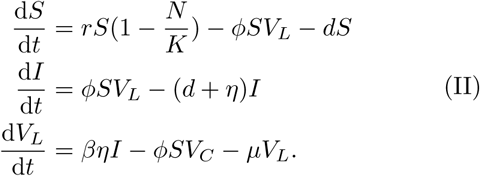

Previous results suggest that lytic virus-host interactions results in three distinct asymptotic outcomes (assuming *r* > *d*): virus clearance, steady state or oscillatory dynamics [23]. Indeed, the lytic subsystem (II) has an infection-free equilibrium,

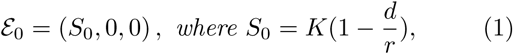

and persistence versus extinction depends upon the lytic infection threshold 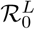:

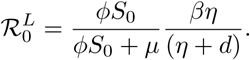

The reproduction number 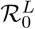 is formulated differently than the threshold derived in [23], but both formulas are equivalent as threshold quantities. Here, the biological interpretation of 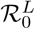 is the average number of secondary cases (infected with lytic viruses) produced by one infected cell (or virus) during its life span in a wholly susceptible microbial cell population. The first term 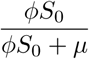 is the probability of a virus infecting a cell before decaying and the second term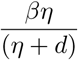 quantifies the average number of new viruses produced by one infected cell in its lifetime. Berretta and Kuang [23] show (in a rescaled version of (II)) the following: If 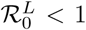, the lytic virus population eventually dies out and the susceptible cell population converges to the equilibrium *S*_0_. Otherwise if 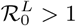, the virus (*uniformly*) persists and the populations converge to the positive equilibrium 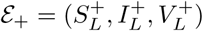, with

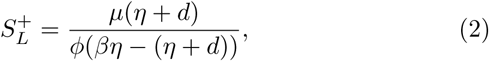

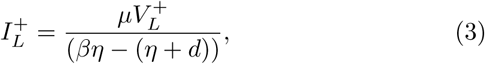

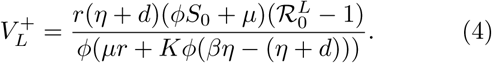

if it is locally asymptotically stable. Yet under certain conditions, this equilibrium 𝓔_+_ loses its stability through Hopf bifurcation, in which case both virus-hosts populations undergo sustained oscillations [23] (see Fig. 1).

**FIG. 1:**
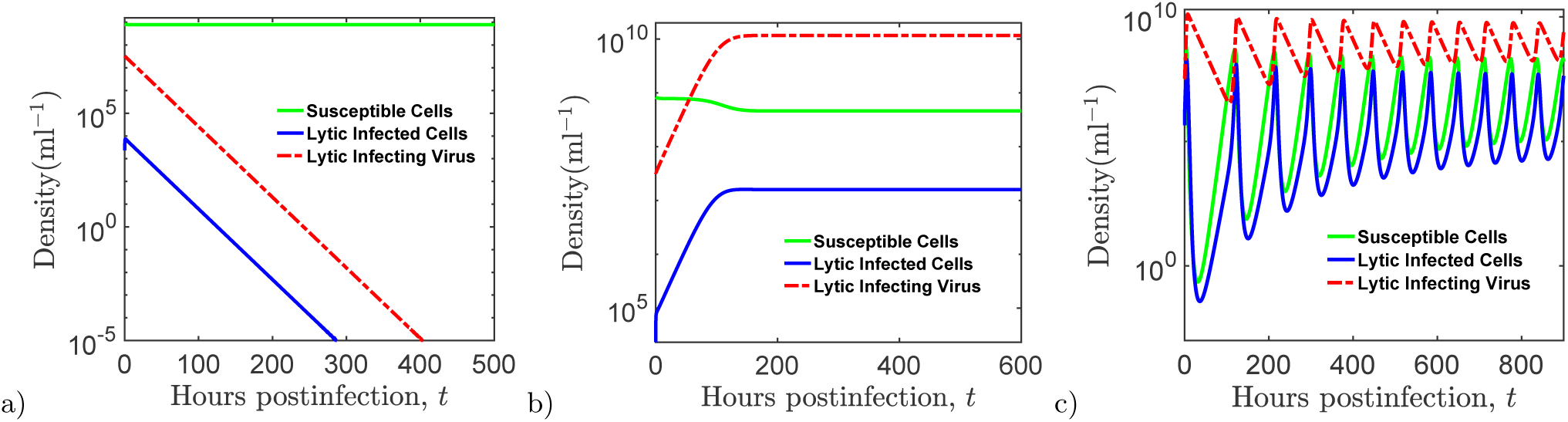
Dynamics of the lytic-subsystem (II) with susceptible host, *S*(*t*), lytic infected host, *I*(*t*), and lytic infecting free virus population density, *V_L_*(*t*), at time *t. a*) Infection dies out and solutions converge to infection-free equilibrium 𝓔_0_. b) Solutions converge to steady-state equilibrium 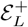. c) Stable positive equilibrium, 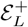, undergoes Hopf bifurcation and solutions present sustainable oscillations, converging to a limit cycle. Common parameters for the dynamics are given in the Table IV. The initial virus and host densities are 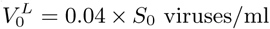, *S*_0_ = 8.3 × 10^8^ hosts/ml. For part (a) *ϕ* = .0.1 × 10^−11^, part (b) *ϕ* = 0.1 × 10^−10^ and part (c) *ϕ* = 0.55 × 10^−9.5^ with *η* = 1.5.

### B. Chronic Infection Outcomes

Next, we consider the subsystem of the model (I), where *V_L_* (*t*) = 0 and *Ι*(*t*) = 0, describing the interactions and the dynamics of hostnumber of chronically infectin gand chronically infecting virus populations:

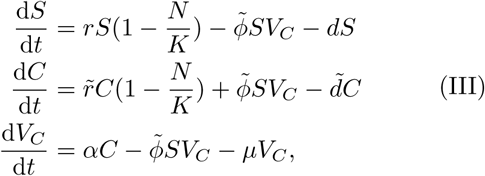

where *N* is the total number of cells, *N* = *S* + *C*. Both lytic-only and chroniconly subsystems have the same infection-free equilibrium, 𝓔_0_, characterized by *S*_0_, the equilibrium level of susceptible cells in the absence of infection. Utilizing the Next Generation Matrix Approach (see Appendix B 1), we obtain the reproduction number of chronically infecting virus 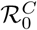:

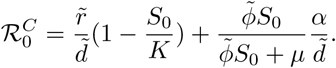

The threshold, 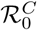, gives the average number of secondary chronically infected cells produced by one chronically infected cell during the life span in a wholly susceptible cell population. The first term 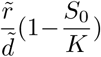 is *the average number of offsprings* produced by an average chronically infected cell during its life span (through vertical transmission) and the second term 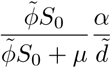 describes the average number of secondary chronically infected cases produced by one chronically infected cell during its life span among completely susceptible cell population (through horizontal transmission). Notice the additional first term compared with the lytic reproduction number 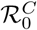. So even in the absence of horizontal chronic infection 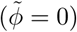, chronic viruses and infected cells can persist so long as:

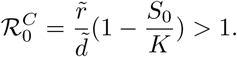

The chronic subsystem (III) exhibits more possible *boundary* equilibria than the lytic subsystem (II). There is the infection-free equilibrium 𝓔_0_ as defined for the lytic subsystem (1). In addition, there can be another boundary equilibrium, namely a chroniconly equilibrium given by

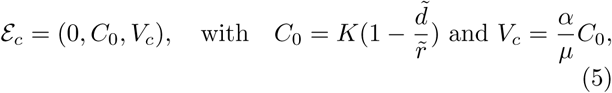

which exists when 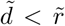. The chronic subsystem (III) also can have one or two positive *interior* equilibria,

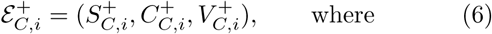

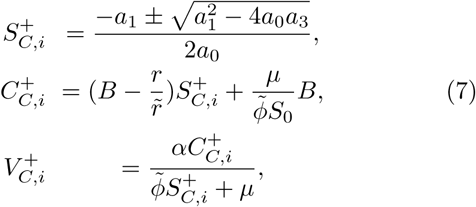

with

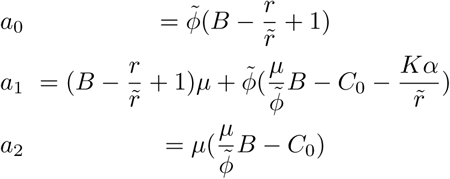

and 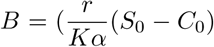 (for derivations, see Appendix C2). Finally, the trivial community collapse equilibrium, 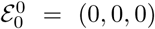, always exists, but is unstable as long as 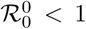, where 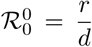. For the rest of the paper, we assume 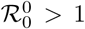, which is a necessary and sufficient condition for existence of the infection-free equilibrium 𝓔_0_ = (*S*_0_, 0, 0).

Given all possible equilibria of the chronic subsystem, the stability analysis of the system (III) provide us crucial information for the complex dynamics displayed by the system (III). First, if 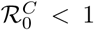, then 𝓔_0_ is locally asymptotically stable (Theorem B.1 in Appendix (B 1)). Yet, this condition does not guarantee extinction of the virus for all initial conditions. In particular, there are parameter regimes with *bistable dynamics* or *bistability*, which generally refers to a dynamical system containing multiple stable equilibria and/or limit cycles with distinct basins of attraction [33]. In other words, the initial virus or infected cell density affects to which attractor the solution converges.

A necessary condition for occurrence of bistability can be analytically expressed as a local condition at 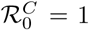. Considering 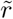 as a bifurcation parameter, backward bifurcation occurs at the critical bifurcation point 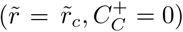, where 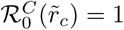, if the following condition satisfies:

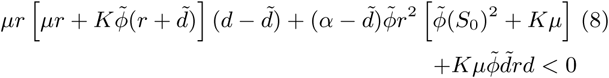

(for derivation, see Appendix (C3)). This condition signals the presence of an unstable positive interior equilibrium when 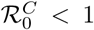, but sufficiently close to one, which will intersect with the infection-free equilibrium 𝓔_0_ at 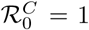, exchange stability and become negative when 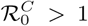. Note that *backward* bifurcation refers to the slope of the positive equilibrium at this transcritical bifurcation. Transcritical bifurcation is a type of bifurcation, where at the critical value of 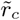, the stability of two equilibria with one stable and one unstable equilibrium, switches as they pass through at this critical point. It is a *backward* bifurcation when the system exhibits an unstable positive interior equilibrium along with the stable infection-free equilibrium 𝓔_0_ (or stable chroniconly equilibrium 𝓔*_c_*). The unstable positive equilibrium (when 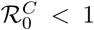), which satisfies one of the equations (6), forms part of a separatix, separating basins of attraction of distinct attractors. One of the attractors is 𝓔_0_, since 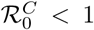 is the condition for local stability. Interestingly, the other attractor can vary depending on parameter region, as model (III) displays multiple types of bistable dynamics.

**TABLE I:**
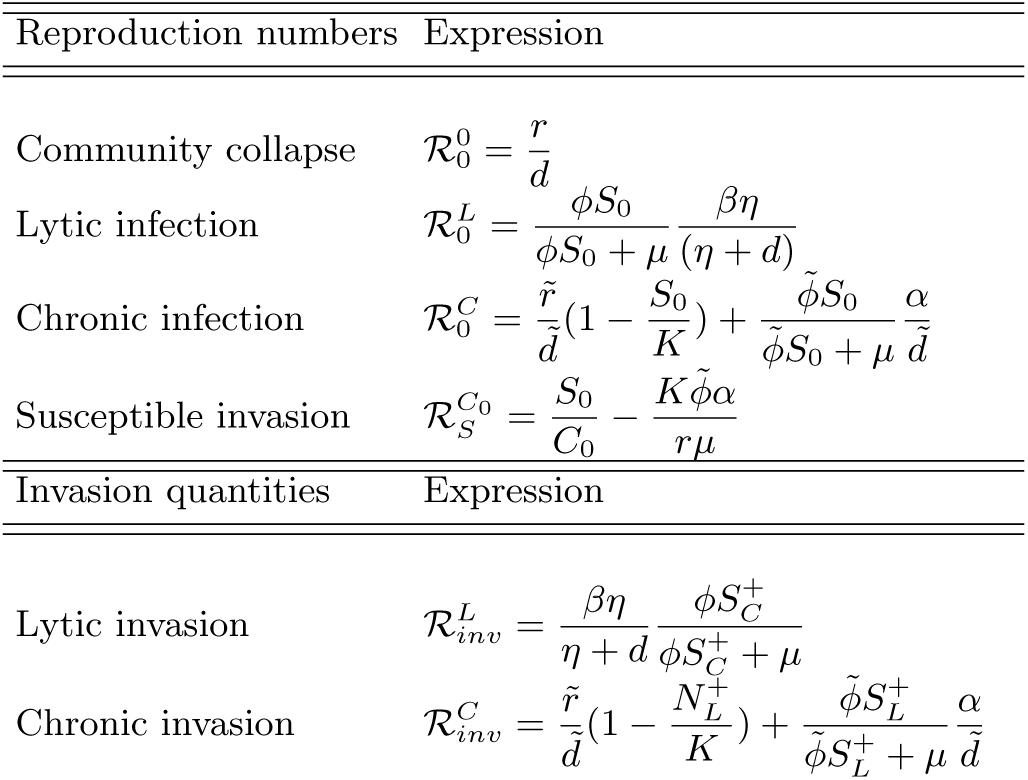
Lytic & Chronic Threshold Quantities Reproduction numbers Expression

**TABLE II:**
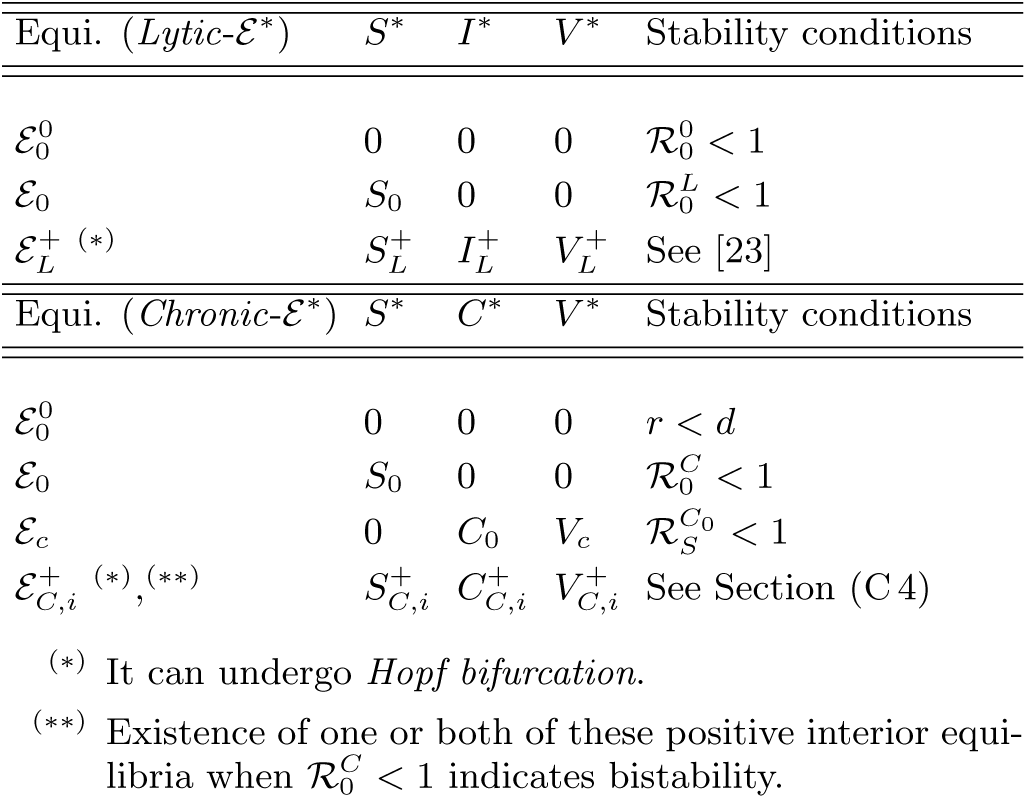
Lytic & Chronic Equilibria & Stability Conditions

One form of bistability in model (III) occurs when both the infection-free equilibrium 𝓔_0_ and the chroniconly equilibrium 𝓔*_c_* are locally stable, and thus are both attractors. The condition for local asymptotic stability of 𝓔*_c_* (Theorem C.2 in Appendix (C 1)) is given by

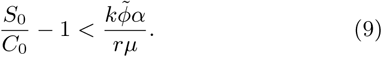

We define the susceptible invasion reproduction number (calculated at the chronic-only equilibrium) as:

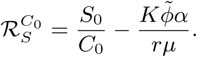

If the threshold quantity 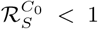, the chroniconly equilibrium 𝓔*_c_* is locally asymptotically stable, and otherwise if 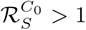, then 𝓔*_c_* is unstable. Again considering 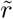 as a bifurcation parameter, there is a transcritical bifurcation point 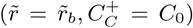, where 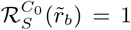. Here the chroniconly equilibrium 𝓔*_c_* exchanges stability with a positive interior equilibrium and the *S* component of this equilibrium, 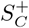, becomes negative. Considering 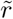 as a bifurcation parameter, backward bifurcation occurs at the critical bifurcation point 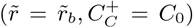, where 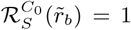, if the following condition satisfies:

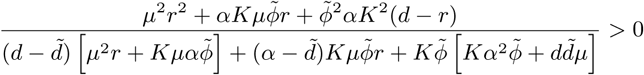

(for derivation, see Theorem C20 in Appendix).

If 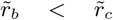, then both transcritical bifurcations at 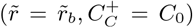, and 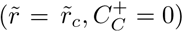 are “backward”. In this case, if 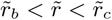, or equivalently 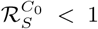 and 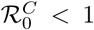, then the fates of the chronic and susceptible host populations depends on the initial size of chronically infected host-virus concentration as shown in Figure (2)(a)-(b). Numerically, we observe that if the initial virus or chronically infected cell concentration is low, then the virus goes extinct; however, for large enough initial concentrations, the system converges to the equilibrium state, 𝓔*_c_*, where only chronically infected cells survive. The biological intuition is that for certain parameter regimes, higher chronically infected cell densities allow the chronically infected cells to outcompete susceptible cells. So in an environment, where chronic cell/virus density is large, even if 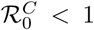, virus can change the fate of the cell population in such a way that chronically infected cells are sustained. We might observe bistability for the case: 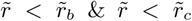 as well, since in the parameter region, where 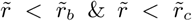, the system has none, one or two positive interior equilibria, 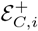 for *i* = 1, 2 (Theorem C.5d in Appendix). The Fig. 3(c) shows the parameter region where this type of bistable dynamics occurs. The corresponding region in the bifurcation diagram, Fig. 3(d), shows that in addition to the stable infection-free equilibrium, 𝓔_0_, and stable chroniconly equilibrium, 𝓔*_c_*, there is one unstable positive equilibrium 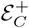 (in certain cases there can be two unstable interior equilibria; see Fig. E.5(h) in Appendix).

**FIG. 2:**
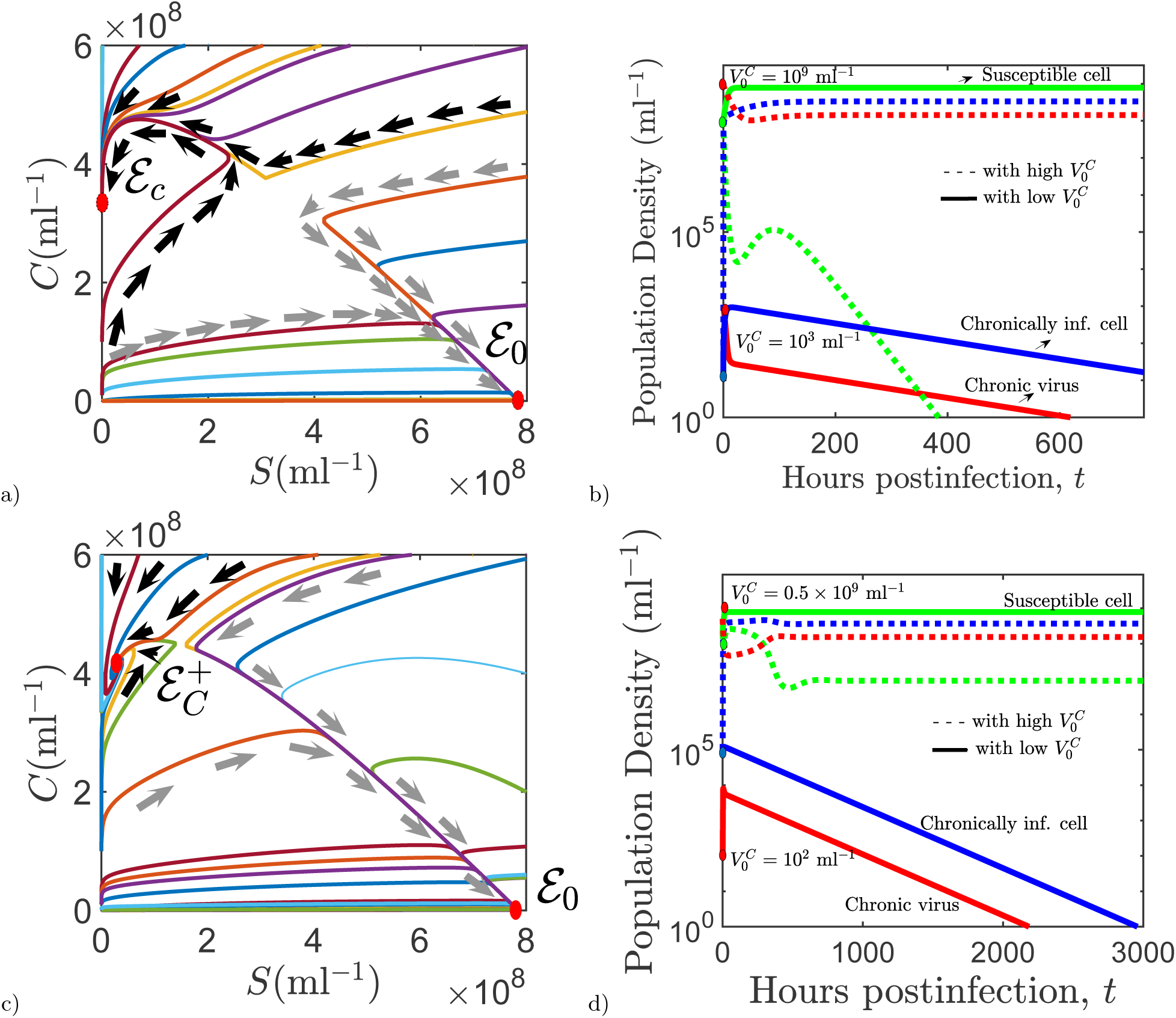
Bistable dynamics displayed by chroniconly system (III). a) Phase plane of the system (III), where bistability occurs with local stable chroniconly equilibrium 𝓔*_c_* = (0, *C*_0_, *V_c_*) and infection-free equilibrium 𝓔_0_ = (*S*_0_, 0, 0). b) Corresponding time dependent solutions of the system (III). Chronically infected cells competitively exclude the susceptible cells when the initial chronic virus density, 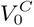, is high or vice versa for low initial virus density size 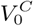. The parameter values are identical to the ones in Table I, except 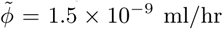, 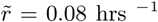, *α* = 1/27. The initial virus and host densities are *S*_0_ = 10^8^ viruses/ml, *C*_0_ = 10 hosts/ml, 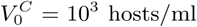 (low density) and *V_C_*(0) = 10^9^ hosts/ml (high density). c) Phase plane of the system (III), where bistability occurs with infection free equilibrium 𝓔_0_ and positive equilibrium 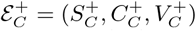, where susceptible and chronically infected cells coexist. d) Corresponding time dependent solutions of the system (III). Bistability occurs with positive equilibrium (with high 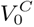) and infection free equilibrium (with low 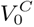). The parameter values are identical to the ones in part (a)-(b), except 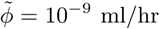. The initial virus and host densities are *S*_0_ = 2 × 10^8^ viruses/ml, *C*_0_ = 10^5^ hosts/ml, 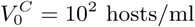 (low density) and 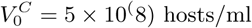 (high density).

**FIG. 3:**
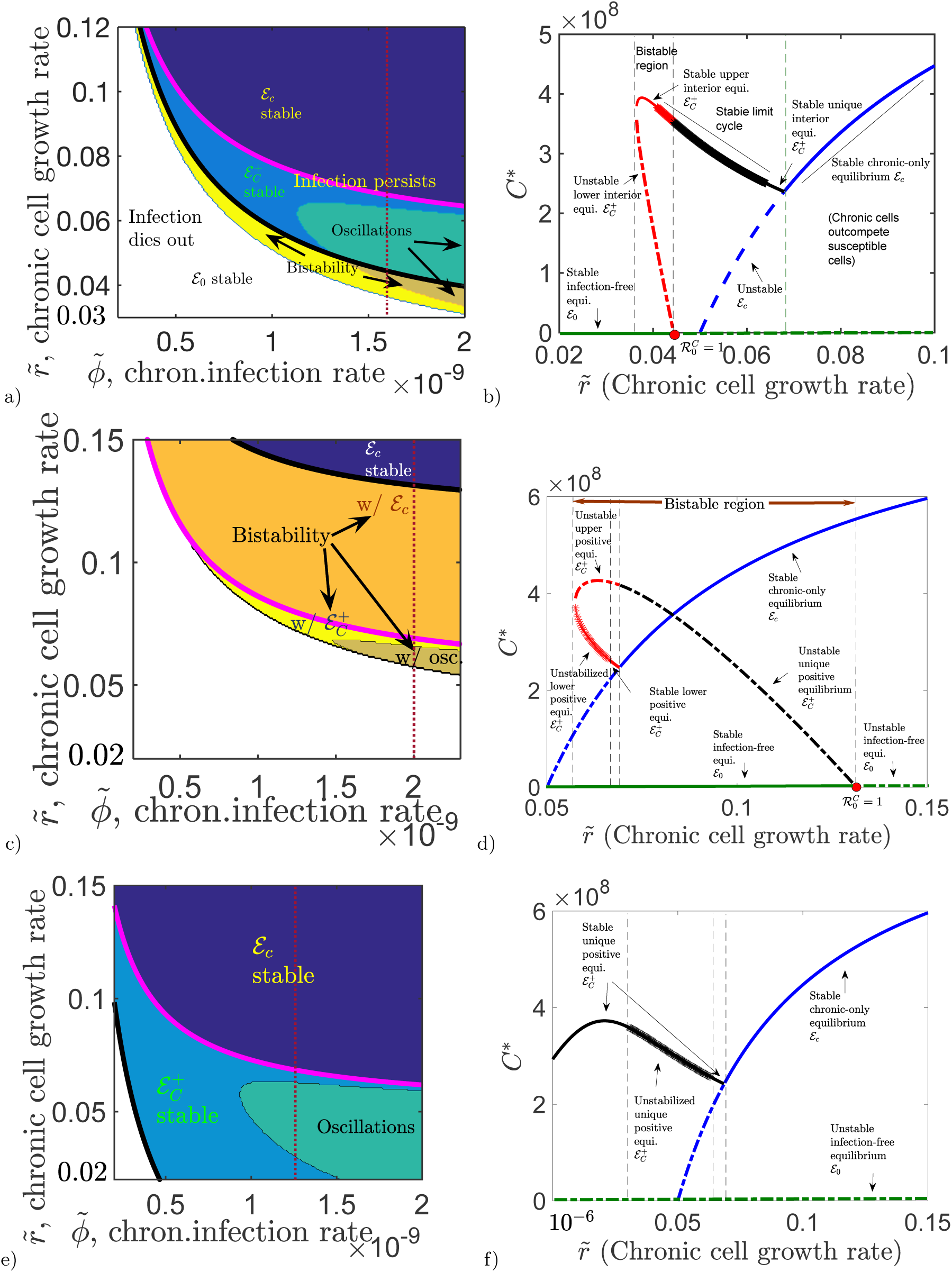
Distinct chronic infection regimes and complex bifurcation dynamics displayed, by varying parameters. a)Distinct chronic infection regimes when *α* = 1/21 b) Corresponding bifurcation diagram of the chronic subsystem (displayed in the parameter region part (a)) when 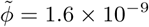 (See the vertical dashed red line). c) Distinct chronic infection regimes when *α* = 1/28, (d) Corresponding bifurcation diagram when 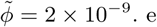) Distinct chronic infection regimes when *α* = 1/17, f) Corresponding bifurcation diagram when 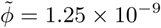. The rest of the parameter values are identical to the ones in Table I. Sustained oscillations, obtained via Hopf bifurcation, is shown with ★.

The other type of bistability consists of the infection-free equilibrium 𝓔 o and a positive interior coexistence equilibrium, 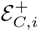, given by (7)) or a positive periodic solution as attractors. In this case, condition (8) holds, along with 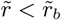 and 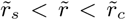, where 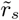 is the location of a saddle-node bifurcation. *Saddle-node* bifurcation is a type of bifurcation, where at a critical value of 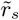, the bifurcation diagram branch out two equilibria with one stable equilibrium and one unstable equilibrium. We observe this type of bifurcation in Fig. 3(b)-(d). Fig. 2(c) depicts the phase plane diagram of this scenario. The corresponding time-dependent solutions of model variables *S*(*t*), *C*(*t*), and *V_C_*(*t*) are shown in Fig.2(d) for multiple initial conditions 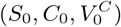. These figures show that larger initial chronic virus concentration 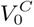 results in coexistence of chronic and susceptible cells, yet when the initial chronic virus concentration, 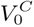, is small, susceptible cells competitively exclude the chronic cell population.

To study the local stability of a positive interior equilibrium, 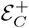, we evaluate the Jacobian Matrix around this equilibrium and study the sign of the real part of the eigenvalues, which are the roots of the characteristic equation derived from the Jacobian Matrix. Analytical and numerical results suggest that in certain cases a positive interior equilibrium can lose its stability via Hopf Bifurcation (section C 4 in Appendix). *Hopf* bifurcation is a bifurcation type, where as 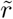 increases (or decreases) the stable interior equilibrium loses its stability at a critical point and displays sustained oscillations. For example, Fig. C.2 in Appendix C 4, shows the interval of 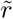, on which the sign of the complex eigenvalues switch its sign from negative (the case 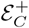 is locally stable) to positive (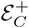 is unstable) or vice-versa. At the critical point, where the real part of the complex eigenvalues become zero, the equilibrium 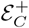 undergoes Hopf bifurcation. We obtain Hopf bifurcation in Fig. 3(b)-(d)-(e) in different settings. In Fig. 3(a), the stable “upper” interior equilibrium undergoes Hopf Bifurcation at a critical value of 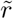 in a bistable region (when 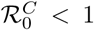). In Fig.3 (a), the system also displays Hopf bifurcation when 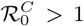 as the unique interior equilibrium stabilizes and limit cycle ceases to exist. Other parameter regimes where Hopf bifurcation of an interior equilibrium occurs are displayed in Fig.3 (d) and (f), including the case where the “lower” interior equilibrium undergoes the Hopf bifurcation.

While there can be two positive interior equilibria when 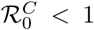, there is at most one positive interior equilibrium when 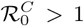. In particular, if 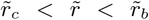 (or equivalently 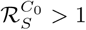 and 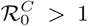), then the system has a unique positive interior equilibrium, 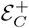 (Theorem C.5 in Appendix). As 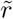 increases above 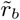, there are no interior equilibria (Theorem C.5(a) in Appendix) and the chronic-only equilibrium 𝓔_*c*_ appears to become globally asymptotically stable. This occurs when 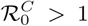 and the 𝓔_*c*_ becomes locally stable 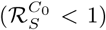. Here, the transcritical bifurcation at 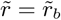 results in the stable interior equilibrium intersecting with 𝓔_*c*_, exchanging stability and becoming negative through the boundary *S* = 0. In addition to that we observe that Fig. 3(b)-(d) exhibits all sort of bifurcation dynamics such as *saddle-node, transcritical* bifurcation (in particular *backward*) and *Hopf* bifurcation as 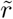 varies.

Distinct bifurcation dynamics displayed by the chronic subsystem (III) are undoubtedly very interesting. We remark that there are even further distinct stability scenarios for equilibria, although they have similar qualitative dynamics to some of the cases described throughout this section (see Fig. E.5 in Appendix.). The different bifurcations depict how the nature of chronic infection mode can significantly change the interactions and population dynamics of viruses and their hosts (see Fig.E.4). As seen in Fig. 3, the varying magnitude of the chronic cell growth rate, 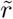, and chronic infection rate, 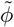, can result in many distinct complex population dynamics outcomes, resulting in significant change in the abundance and the dynamics of host cells and their viruses.

## IV. MULTI-STRAIN MODEL: INVASION DYNAMICS

Next, we move to the full multi-strain model (I) in order to investigate *how interaction of lytic and chronic viruses with a common microbial host can result in distinct ecological outcomes.* Competition between two species or population variants results in competitive exclusion or coexistence. Two species often cannot occupy the same niche, with the more fit species driving the other to extinction. However, heterogeneous strategies can allow for two species to both exploit a common resource and coexistence becomes a possible outcome. The starting point for analyzing competition between two species is to determine *when each species can establish their population in the presence of the other resident species.* In this section, we study under what conditions the lytic (chronic) virus type can successfully invade a resident chronic (lytic) virus population.

First, we formulate invasion fitness quantities of both virus types. An *invasion fitness quantity* is a threshold value, allowing us to infer whether a virus type can invade a distinct resident virus population.

Assuming that by the time at which lytic virus is introduced, the resident chronic virus population is at its equilibrium 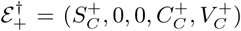, where positive components are given by (7), we obtain the lytic invasion fitness quantity 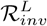 as follows:

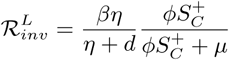

It can be interpreted as the reproduction number of lytic cells at the boundary equilibrium 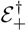 analogous to the basic reproduction number 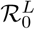, which is calculated at the infection-free equilibrium 𝓔_*0*_ instead.

Analytical results suggest that a subpopulation of lytic viruses invade the chronic resident population if 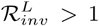.

In more mathematical terms, the boundary chronic infection equilibrium, 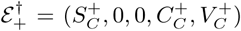, is unstable with respect to invasion if 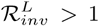 (see Appendix D). If 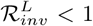, then 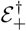 is locally asymptotically stable when the chronic-infection equilibrium 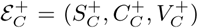 is stable in the chronic subsystem (III). Note that in the parameter region where the resident chronic population is oscillating (at a stable limit cycle) upon arrival of lytic viruses, then the invasion depends on a linear periodic system (given by the Next-Generation matrix at the limit cycle) [36].

We can also formulate the threshold quantity providing whether chronic cells or viruses can invade the resident lytic population, when at the equilibrium 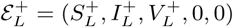, where positive components are given by (2). The *chronic invasion fitness quantity*, 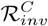, is defined as

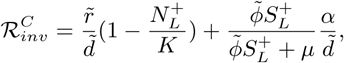

where 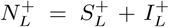. When a resident lytic type is at its equilibrium 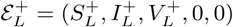, rare (a small initial density of) chronically infecting viruses can only invade if 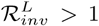. Otherwise if 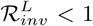, analogous conclusions hold as in the case of lytic invading chronic.

Having both invasion threshold quantities, 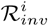, along with basic reproduction numbers, 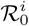, where *i* ∈ {*L*, *C*}, we can study distinct under what conditions invasion of one type or infection take place. Table V in Appendix gives distinct cell and virus fates under distinct values of these invasion and infection thresholds when the initial chronic virus or cell density is sufficiently low. Fig.4 shows distinct parameter regimes where outcomes of interactions varies such that chronic virus can substitute the lytic viruses (R.VI), while fails to invade in another region (R.IX). Chronic invasion can also result in coexistence (R.IV & R.VIII) and exclusion of both types (R.V), which will be further discussed in the next section.

**FIG. 4:**
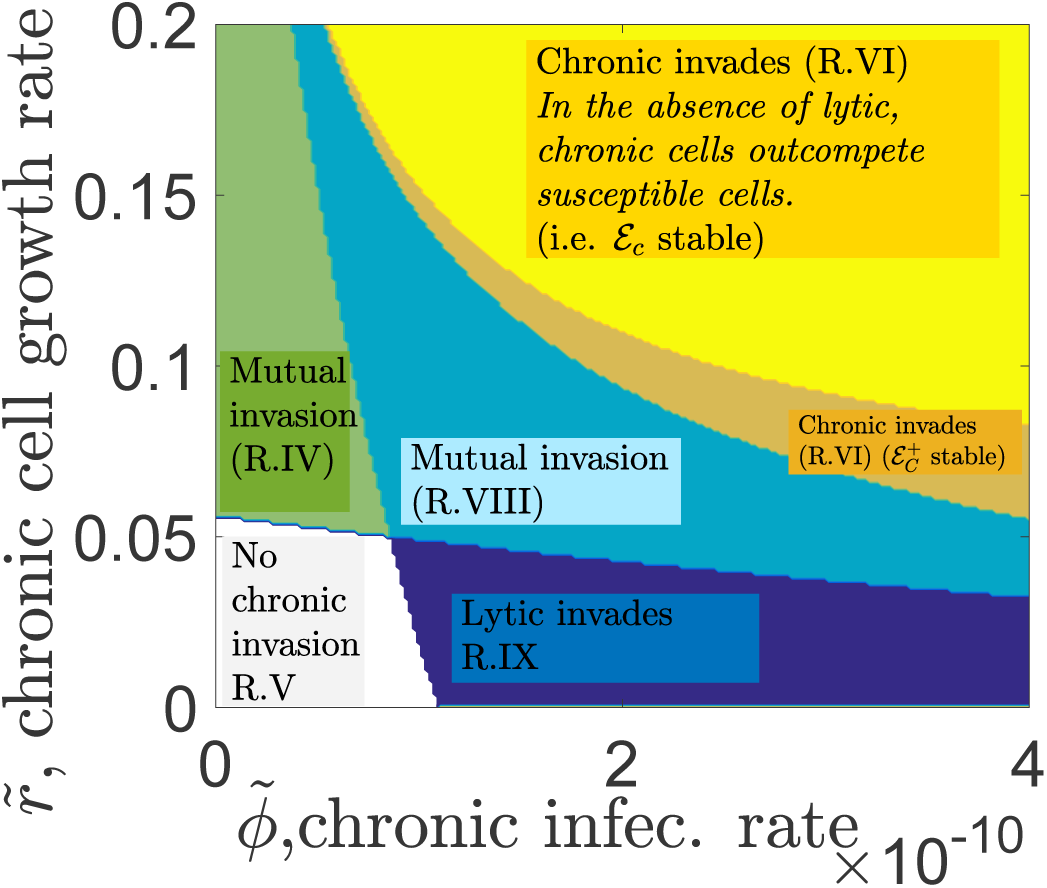
Infection, invasion and mutual invasion parameter regimes of chronically & lytic infecting viruses. On the x-axis, we vary the chronic infection rate, 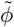, and on the y-axis, we vary chronic cell growth rate, 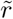. Given the lytic parameter values on this map, we have 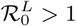. The contour map displays in what parameter regime chronically (lytic) infecting virus invade lytic (chronic) virus population and when it fails to do so. Mutual invasion regimes provide the coexistence region. The parameter values are identical to the ones in Table I. The analytical conditions, providing these regimes, are given in Table V.

## V. COMPETITION & COEXISTENCE OF LYTIC AND CHRONIC VIRUSES

### A. Heterogeneous viral strategies promote coexistence

The multi-strain system (I) can have a unique coexistence equilibrium 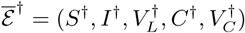, derived as

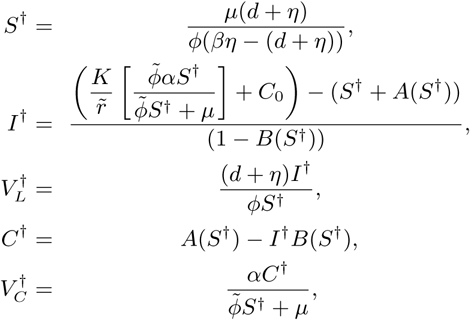

where

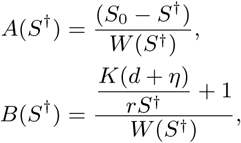

with

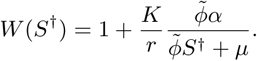

(see Appendix E). The expressions for 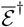 are too complicated in order to analytically determine the conditions for its positivity. However, numerical results show that equilibrium 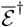 can be positive in certain parameter regions, in particular when both invasion conditions 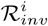 *i* ∈ {*L*, *C*} are greater than one (discussed further below). Therefore, the system can have a unique coexistence equilibrium for certain parameter regimes. In Figure (5), solutions tend to this coexistence equilibrium asymptotically. The coexistence equilibrium 𝓔^†^ can lose its stability via Hopf bifurcation, in which case both virus densities oscillate and converge to a limit cycle as shown in Figure 5(d).

**FIG. 5:**
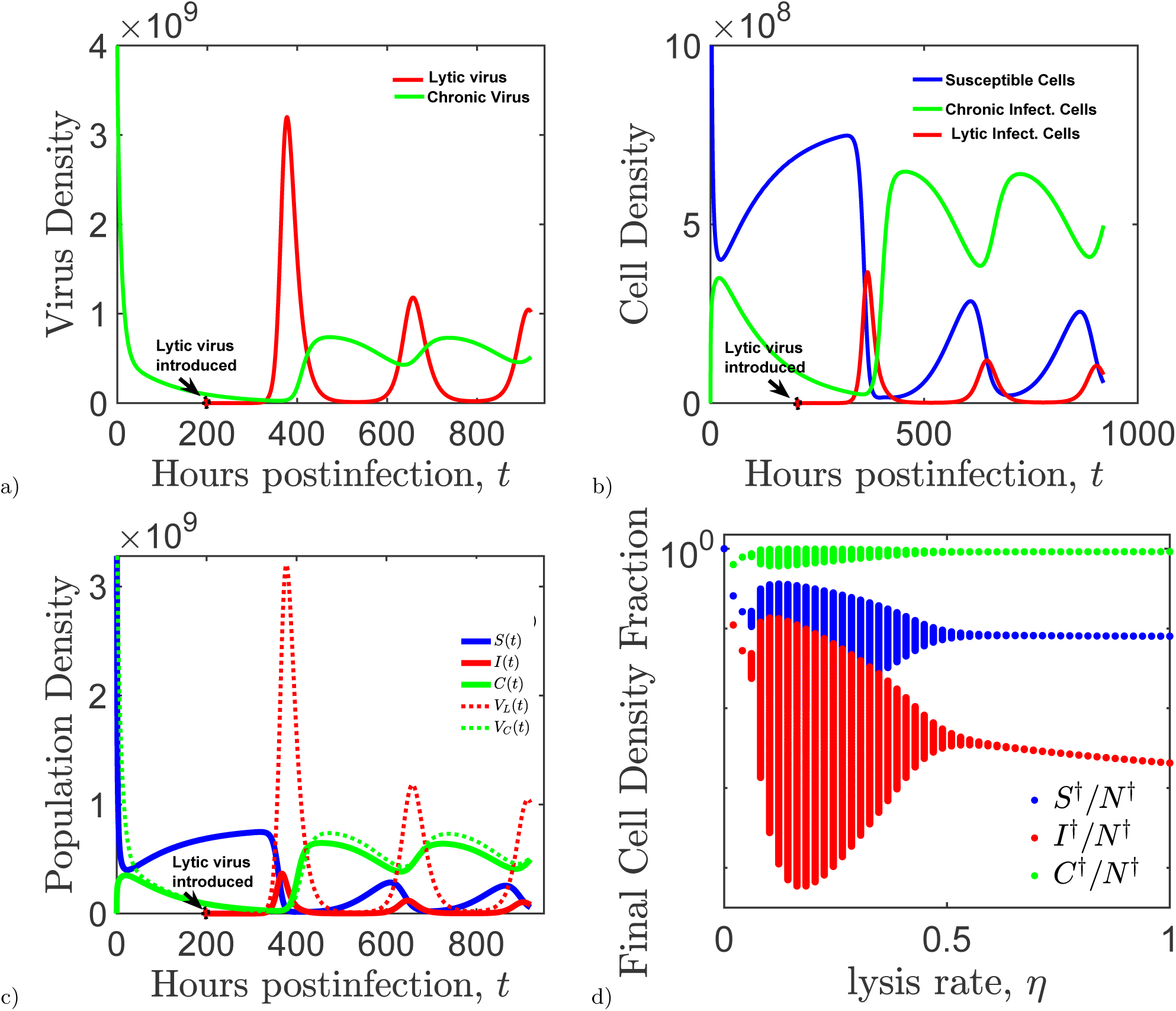
The dynamics displayed, by chronic virus before and after lytic virus introduced. a) “Rescue” of chronically infecting viruses by lytic viruses (*η* = 0.0526). b)Changing microbial host cell population dynamics after lytic virus introduction. b)Microbial host-virus population dynamics before and after introduction of lytic viruses. The initial virus and host densities are *S*_0_ = 10^1^0 viruses/ml, 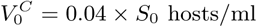, 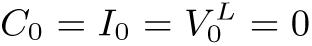 hosts/ml. d) Fraction of asymptotic cell density with respect to virulence rate *η*. The solid vertical lines display the sustained limit cycles with their magnitudes. The parameter values are identical to the ones in Table I.

There are parameter regions where both invasion conditions 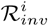 *i* ∈ {*L*, *C*} are greater than one, which suggests persistence of both lytic and chronic virus for these parameters. There are two distinct scenarios where 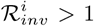, *i* ∈ {*L*, *C*} as shown in Table V. In particular, Region *IV* depicts an interesting scenario where 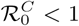 but 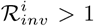, *i* ∈ {*L*, *C*} so that chronic can be wiped out in the absence of lytic virus, yet both viruses persist together in the multi-strain model (this scenario is discussed further in Section VB). The other case of coexistence is where all thresholds 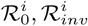 are greater than one (see Fig. 4).

In either case of coexistence, the heterogeneity of the viral strategies is critical for the persistence of both virus strains. Indeed if 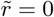, i.e. the chronic virus only reproduces through standard viral replication (budding or bursting of infected cells, as in the case of lytic virus), then there is no coexistence equilibrium except in the case 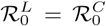. In addition, when 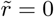, the invasion fitness quantities reduce to 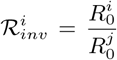 *i* ≠ *j* (shown in Appendix E.3); thus it is not possible for both invasion conditions to be greater than one. Therefore, in our study coexistence of the two virus types hinges upon the additional replication technique displayed by the chronic virus; namely being long-lived and passing on to the microbial host daughter cells after cell division.

**TABLE III:** Multi-strain Equilibria & Stability Conditions

| $Multi\text{-}strain\text{-}\bar{\mathcal{E}}$ | $S^\dagger$ | $I^\dagger$ | $V_I^\dagger$ | $C^\dagger$ | $V_C^\dagger$ | Stability conditions |
| --- | --- | --- | --- | --- | --- | --- |
| $\bar{\mathcal{E}}_0^0$ | 0 | 0 | 0 | 0 | 0 | $\mathcal{R}_0^0 < 1$ |
| $\bar{\mathcal{E}}_0$ | $S_0$ | 0 | 0 | 0 | 0 | $\mathcal{R}_0^i < 1, i \in \{L, C\}$ |
| $\bar{\mathcal{E}}_L^+ (*)$ | $S_L^+$ | $I_L^+$ | $V_L^+$ | 0 | 0 | $\mathcal{E}_L^+$ stable, & $\mathcal{R}_{inv}^C < 1$ |
| $\bar{\mathcal{E}}_c$ | 0 | 0 | 0 | $C_0$ | $V_c$ | $\mathcal{E}_c$ stable |
| $\bar{\mathcal{E}}_{C,i}^+ (*), (**)$ | $S_{C,i}^+$ | 0 | 0 | $C_{C,i}^+$ | $V_{C,i}^+$ | $\mathcal{E}_{C,i}^+$ stable, & $\mathcal{R}_{inv}^L < 1$ |
| $\bar{\mathcal{E}}^\dagger (*)$ | $S^\dagger$ | $I^\dagger$ | $V_I^\dagger$ | $C^\dagger$ | $V_C^\dagger$ | See Section (E2) |

(\*) It can undergo *Hopf bifurcation*.
(\*\*) Existence of one or both of these positive interior equilibria when $\mathcal{R}_0^C < 1$ indicates bistability.

### B. Lytic Virus Facilitates Persistence of Chronic Virus

Lytic infection can alter the dynamics between the distinct reproducing cell types (chronic and susceptible) by reducing cell competition through lysing susceptible cells, thereby facilitating the persistence of chronically infected cells. However this potential (indirect) beneficial interaction goes in conflict with the competition between virus types for common microbial hosts.

How the lytic and chronic virus interactions modulate the biodiversity of the virus-microbe system depends on the parameter regimes upon which the fitness quantities change. *A dramatic instance of lytic virus benefiting chronic virus is in the regime IV*, *where the chronic virus requires the lytic virus for survival and invasion*. For this case, the chronic virus population will not become established in the absence of lytic virus, yet if lytic virus is present in the system, then the chronic virus will persist (See Fig.5). The biological reasoning behind this unexpected result is the following: In a wholly susceptible population without lytic virus, chronically infecting cells face higher cell competition due to larger total microbial cell density, whereas if lytic virulent viruses are present, the amount of cells is decreased through lysis, reducing cell competition, which increases net reproduction of chronically infected cells, allowing for chronic invasion (See Fig.6). Indeed, analytically we observe that a virulent lytic virus type reduces total cell population 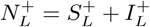, which increases the average chronic offsprings produced, 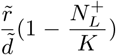, above what it would be in the absence of infection. In particular, for parameter values in regime *IV*, we obtain 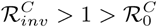 (see Fig.4b).

**FIG. 6:**
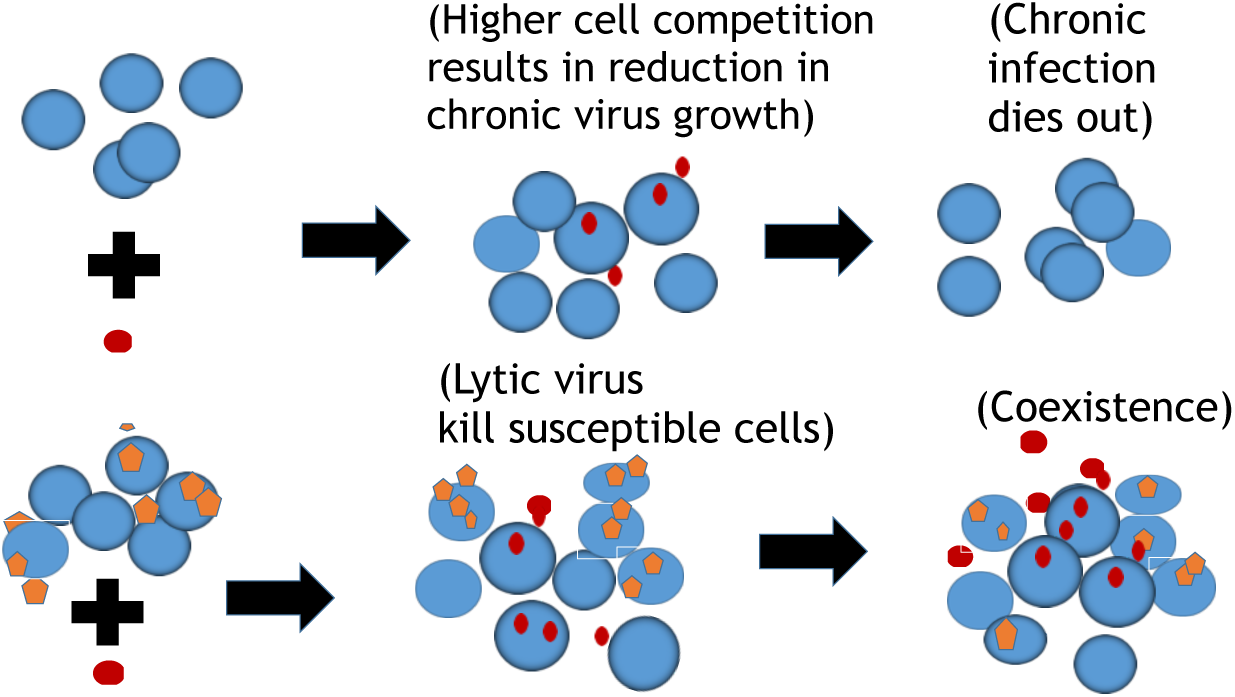
Schematic representation of the outcome of interactions when a rare chronic virus (red) introduced in the absence (upper fig.) or existence of a lytic infecting virus (orange) population (lower fig.). The chronic infection can persist only when lytic infection is already established among the susceptible host population.

We also notice that this beneficial interaction only goes one way as lytic infection can not be “rescued” by chronic viruses. Lytic viruses are more apt to persist without chronic infection since they can access a larger amount of susceptible cells (higher 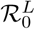), and there is no reverse effect of cell competition on their growth. Analytically, we see that

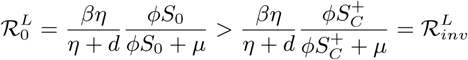

since *S*_o_ is always greater than 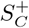.

It is *interesting* that in the regime *IV*, chronically infected cells not only are rescued from extinction by the lytic virus, the chronically infected population can persist at the top of the cell abundance hierarchy (infected or susceptible). Indeed, observe Fig.5(d), where we show how the final size of abundance of susceptible, chronically and lytic infected host population at their equilibrium changes with varying lysis rate, η. If lytic virus were removed from the system or become extinct due to sensitivity to stochastic fluctuations, susceptible cells replenish again and as a result of host competition, the chronically infected cells would face extinction again. So chronically infected cells benefit from the mutual relationship with lytic virus. This coexistence mechanism which favors survival and large chronically infecting virus abundance may explain both the observed virus diversity and could suggest a means to explain findings of intracellular persistence of viruses in natural systems.

## VI. CONCLUSIONS

Virus-microbe interactions are characterized by distinct adaptations, infection modes, and life cycles. Understanding microbial and virus evolutionary strategies requires an understanding of how infection modes couple to population and evolutionary dynamics. In this study, we introduced a model of distinct viral infection strategies (lytic & chronic) interacting with a common microbial host population. In doing so we find that chronic virus infections affect microbial host dynamics, and can induce more complex dynamics than in the case of lytic infections. In addition, in environments where lytic and chronic viruses compete, the heterogeneity in their infection modes promotes the coexistence of the two virus species. Surprisingly, we find that the presence of lytic viruses can benefit chronic viruses, even causing the persistence of chronic infections that would not otherwise be sustainable. Finally, we observe that enhanced virulence of lytic viruses can be more beneficial for persistence chronic infection, which in turn can be detrimental for the lytic virus abundance.

The enhancement of chronic infections due to viral infections warrants elaboration. Our analysis suggests that lytic viruses directly modulate niche competition amongst cells. Hence, inclusion of lytic viruses can decrease susceptible cell populations which, in turn, decreases competition with cells infected by a chronic viruses. Because part of the fitness of chronically infecting viruses derives from vertical reproduction, then lytic viruses can increase chronic virus fitness. The consequences of this phenomena are evident in those examples in which a chronic virus population cannot survive in the absence of lytic viruses. In essence, *lytic modulated cell competition* changes the fate of cell and virus populations in a complex community.

As was noted, chronic infection can have substantial effects on the dynamics of microbial host abundances. In certain parameter regions, chronic infections give rise to similar dynamics to lytic infections: when 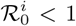, *i* ∈ {*L*, *C*}, the virus population dies out; otherwise if 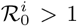, *i* ∈ {*L*, *C*}, both virus and cell populations persists at an endemic equilibrium or at a sustained oscillation, when the positive equilibrium loses its stability via a Hopf Bifurcation. Yet, the fate of chronically infecting viruses or infected hosts might also depend on the initial chronic virus density; i.e. the system presents bistable dynamics, where the initial conditions alter ecological trajectories of both microbial hosts and their viruses. Here, the system contains multiple attractors: the infection-free equilibrium and either a positive interior equilibrium, a positive periodic solution or the boundary chronic-only equilibrium. In each bistable case, chronic viruses can persist even when their reproduction number is less than one, in contrast with lytic viruses.

The results of our analysis raises a question: what is the expected evolution of quantitative traits associated with each infection mode? In the virus-microbe systems we have analyzed, increasing virulence (*η*) decreases the life span of infected microbial hosts (1/*η*), which may in return decrease the amount of virus particles releases during lysis because of physiological limits, but here we assume the burst size *β* remains constant. Our results suggest that the evolution of virulence of lytic viruses can be influenced by the competing chronic virus species. Numerically and analytically, we observe that lytic virus abundance drastically decreases upon the lysis rate *η* increasing past the critical value where chronically infection virus invade and coexist (see Fig.7). Conversely, less virulent viruses can draw the chronic virus to the extinction. Thus in an environment, where susceptible cells can sustain themselves in high abundance, the lytic virus might evolve toward being less virulent, thereby outcompeting the chronic virus population and persisting at much higher abundance. This hypothesis may be suitable for future work, e.g,. extending these arguments by studying evolutionarily stable strategies in this mode or in an extended version with mixed strategies, i.e. one type virus display both type infection modes.

**FIG. 7:**
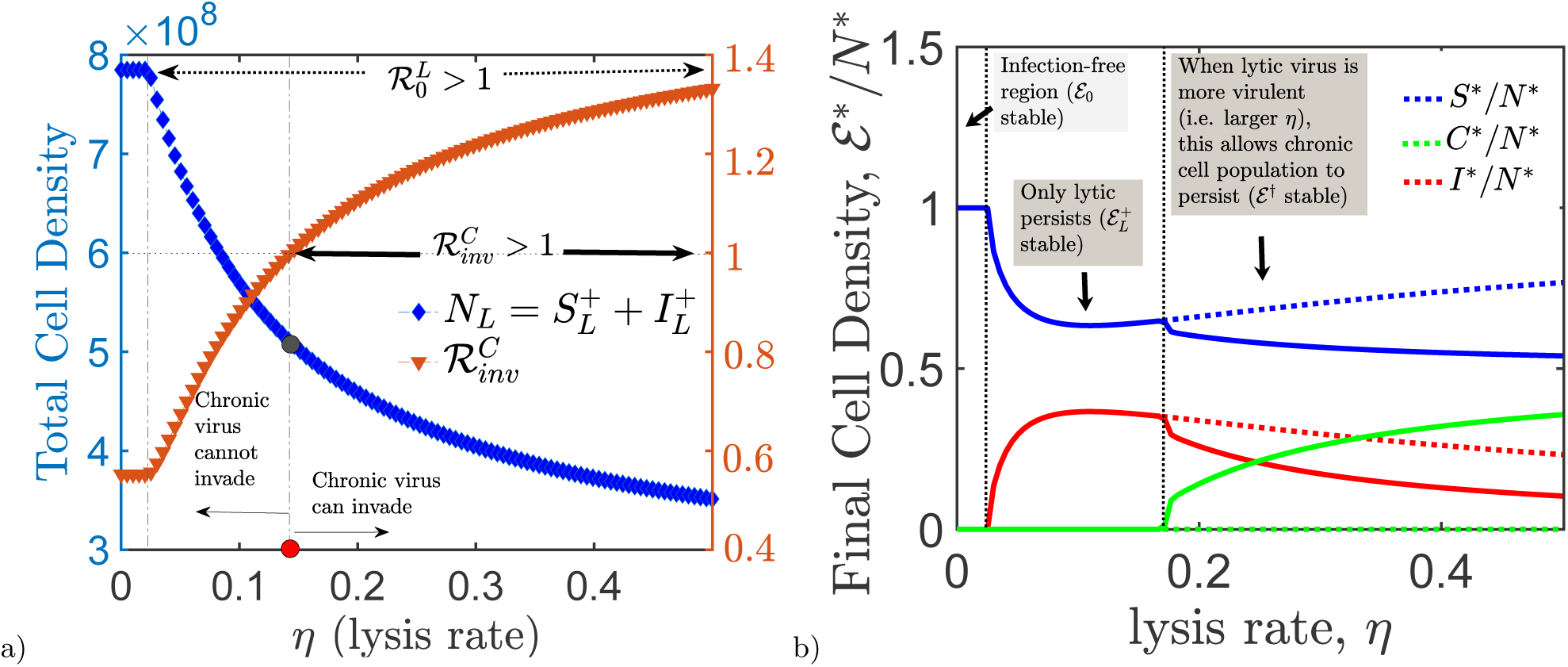
How virulent lytic virus effects invasiveness of chronic virus. a) *Invasiveness of chronic virus vs. lytic virus virulence.* As lytic virus become more virulent, it decreases the total cell density, 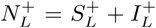, at the equilibrium and increases the invasion fitness, 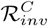, of chronic virus. If 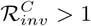, chronic can invade the lytic resident population; otherwise, if 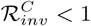, it dies out. b) *The final cell density fraction with respect to varying lysis rate η.* The dashed line represents the fraction of lytic and susceptible cell densities at steady state in the absence of chronic infection. The solid lines display the fraction of cells densities at the equilibrium after chronic infection is introduced. The chronic infection can only persist, when lytic virulence, η, is large enough. The parameter values are identical to the ones in Table I, except 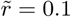, *ϕ* = 0.25 × 10^−10^ and *β* = 15.

There are many ways by which virus interactions can affect the ecology and evolution of virus-microbe systems [3]. Distinct physiological states, life strategies and adaptations emerge in response to the frequent interactions between viruses and microbes. For example, bacteria have evolved adaptive immunity[10], and may also display bet-hedging heterogeneous susceptibility to viruses[15, 16]. Viruses can also exhibit distinct strategies as they interact with microbial hosts[21, 24]. The present study has examined the consequences of heterogeneous viral strategies by analyzing a model of competition between lytic and chronic modes of infection. The present model may also serve as a suitable basis for extensions to cases where viruses display mixed strategies or to the study of the evolution of infection-associated traits. If we are to predict host and virus evolutionary trajectories, it is important to first understand the complex interactions, individual population dynamics and the outcomes of interactions in an ecological setting. Future work linking ecological dynamics and evolutionary trajectories will be necessary to improve our understanding of the complexity and structure of virus-microbe systems.

## Appendix A Parameter Estimations

Gulbudak and Weitz [16] estimate the virus absorption rate *ϕ* = 2.2 x 10^−9^ ml/hr from a recent experimental study of Bautista et al [15], in which the interactions takes place between the archaeon *Sulfolobus islandicus* and the dsDNA fusellovirus *Sulfulobus* spindle shaped virus (SSV9). S. *islandicus* is a globally distributed archaeon, commonly found in hot spring ecosystems. Host growth rate and carrying capacity in the absence of viruses are also estimated as *r* = 0.3 hrs^−1^ and *K* = 9 × 10^8^ cells/ml, respectively. Here we consider these parameters values to be *r* = 0.339 hrs^−1^, K = 8.947 x 10^8^ cells/ml, and *ϕ* = 0.88 × 10^−10^ ml/hr. The chronic virus absorption rate is also fixed as 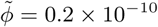. The cell decay rate, *d,* is estimated as *d* = 1/24 hrs^−1^ in [16]. Despite the fact that the chronic infection is not virulent, it might reduce the life span of chronically infected cells due to the infection cost. Hence we fixed the value of chronic cell death rate, 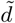 to be 1/20 hrs^−1^. Similarly, virus decay rate is estimated from free virus data in [15] as *μ* = 0.0866 hrs^−1^. In addition,Beretta and Kuang [23] estimates virus replication factor number in the range of 10 to 100 mature virus particles per day. Assuming that not all viral particles produced are infectious, we consider *β* to be the effective burst size and fixed as *β* = 20. Beretta and Kuang [23] also estimate the lysis rate, *η* to be *η* = 3.3/24 (≈ 0.138) hrs^−1^. Here we fixed this value as *η* = 0.33. Although the value of chronic infection parameters, 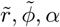, α have varied, through this study, for multiple simulations, when not varied, we chose the value of chronic cell growth rate to be 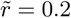, smaller than susceptible growth rate *r*, and the chronic cell virus budding off rate to be α = 1/10. We can find *the average number of infectious viruses produced by one chronically infected cell* by multiplying the budding off rate α with cell division doubling time *τ,* which can be estimated by 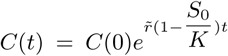. This estimation gives us the average number of infectious viruses produced by one chronically infected cell as ατ ≈ 2.8.

## Appendix B Chronic Infection Dynamics Analysis

### 1. Finding 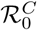 by using Next-Generation approach

The system has an infection-free equilibrium 𝓔_0_ = (*S*_0_,0, 0) with 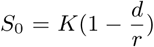. Let the entries of the matrix 𝓕 be the rates of appearance of new chronic infections, and the entries of the transition matrix 𝓥 be the rates of transfer of individuals into or out of compartments such as death, infection, or absorption. Then the Jacobian matrix 𝓘 evaluated at the infection-free equilibrium 𝓔_0_ = *(S*_0_, 0, 0) is 𝓘_𝓔o_ = (𝓕 – 𝓥)|_𝓔o_:

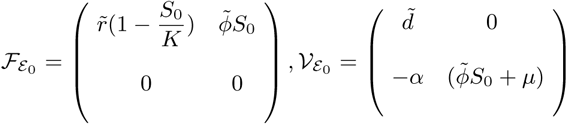

Then by Next Generation Matrix approach [38–40], the spectral radius of Next Generation Matrix 𝓕𝓥^−1^|_𝓔o_ gives *basic chronic reproduction number,* 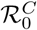:

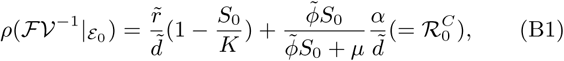

establishing the following theorem:

#### Theorem B.1

(Local stability of 𝓔_0_) Consider the infection transmission model, given by (III). Then if 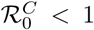, the infection-free equilibrium, 𝓔_0_, is locally asymptotically stable, but unstable if 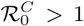, where 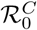 is defined by (B1).

## Appendix C Preliminaries of Chronic Infection Model

To simplify the system (III), we first use the dimensionless time, 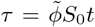, and then rescale the variables of the model (III) by letting 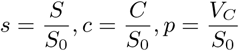. Therefore we obtain the following system:

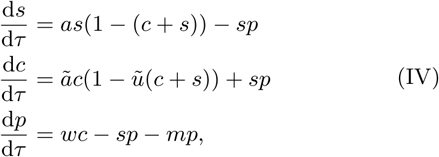

where 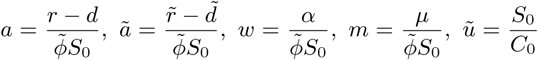
with 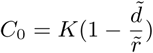

### 1. Existence and stability of chronic-only equilibrium

Let *(s*,c*,p*)* be the equilibrium of the system (IV). Assuming *s*\* = 0, by the equilibrium conditions, we obtain 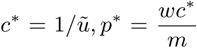. This establishes the following result for the original system (III):

#### Theorem C.1

*The chronic subsystem (III) always has the chronic-only equilibrium 𝓔_c_ =* (0, *C*_0_,*V_c_*), *where* 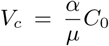 *(recall that* 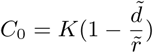).

#### Theorem C.2

*The chronic-only equilibrium, 𝓔_c_, is locally asymptotically stable if and only if the condition* 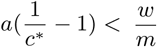 *holds which is equivalent to the condition*:

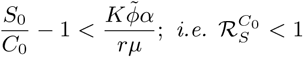

*in the non-dimensionalized original system (III). Otherwise,* 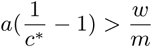, *(or* 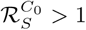*), then 𝓔_c_ is unstable*.

***Proof C.1** By linearizing the system (IV) around the equilibrium 𝓔_c_ = (0, c*,p*), we obtain the following characteristic equation:*

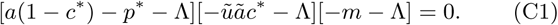

*Then the Jacobian Matrix of the system (III) evaluated at 𝓔_c_ = (0, c*,p*), has all eigenvalues, Λ, negative if the condition* 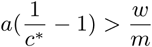 *holds. Otherwise, if* 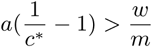, *it has one positive eigenvalue*.

Furthermore assuming both conditions 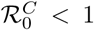 and 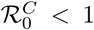 hold, then the fates of the chronic and susceptible host populations depend on the initial chronically infected host and virus concentration; i.e. the system exhibits bistable dynamics. There are cases, where bistabiliy occurs with two interior equilibria (one stable and another unstable) when 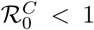. We will derive the *bistability condition* for general case in Section (C 3).

### 2. Existence of Positive Equilibria

Here, we define positive equilibrium to be the equilibrium (*s*,c*,p\**) with all components in the positive orthant. By the equilibrium conditions, derived from the system (IV), we have

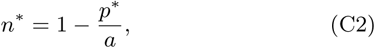

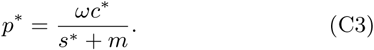

and

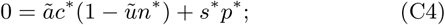

where *n*\* = *s*\* + *c*\*. Substituting both equations (C2) and (C3) into the equation (C4), we get

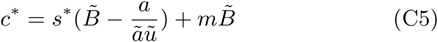

where 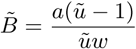. Then

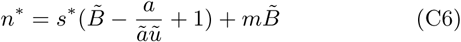

By (C4), we also have

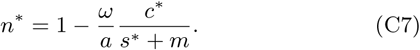

Therefore substituting (C5) into (C7), we obtain

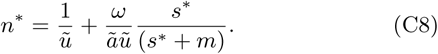

By the equality of the equations (C6) and (C8), we have

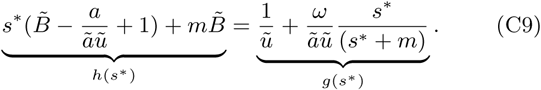

Let the left hand side of the equation be *h*(*s*\*) and the right hand side of the equation be *g*(*s*\*). Both functions *h*(*s*\*), *g*(*s*\*) are monotone, where *h*(*s*\*) is linear function with slope 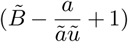 and *g*(*s*\*) is an increasing saturating function of *s*\*, converging to 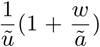 as *s* * → ∞. Recall that we are only considering *s* * ∈ (0,1).

*Existence and number of the positive equilibria depends on the parameter region*:

- Assuming 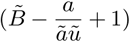, then *h(s*\*) is a decreasing function of *s* *. In which case, if *h*(0) < *g*(0), then the functions *h*(*s*\*) and *g*(*s*\*) do not intersect at a positive value *s* *. Therefore if *h*(0) < *g*(0), then the system does not have a positive equilibrium. Otherwise, if *h*(0) > *g*(0), the graphs *h*(*s*\*) and *g*(*s*\*) intersect at a positive value, namely 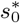 in (0,1), whenever *h*(1) < *g*(1). In this case, the system has a positive equilibrium 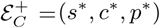 only if 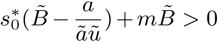. Notice that to obtain a positive equilibrium, we need to also obtain a corresponding positive value for 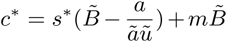 for the positive intersection *s*\*. In summary, given the condition that 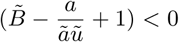, the system can have at most one positive equilibrium 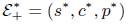.
- If 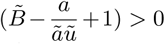, then *h(s*\*) is an increasing function of *s*\*. Thus

- If *h*(0) < *g*(0), then *h* and *g* have one positive intersection *s*_0_ in (0,1) whenever *h*(1) > *g*(1). In which case whenever *h*(0) < *g*(0) and *h*(1) > *g*(1), the system has one positive equilibrium, assuming 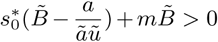.
- If *h*(0) > *g*(0) and *h*(0) ≈ *g*(0), then

* zero equilibrium when *h*′(0) > *g*′(0),
* two equilibrium when *h*′(0) < *g*′(0), and *h*(1) > *g*(1), assuming there is no *s*\* ∈ (0,1) : *h*′(0) = *g*′(*s*\*).
* or one equilibrium if ∃*s*\* ∈ (0,1) : *h*′(*s*\*) = *g*′(*s*\*).
- If 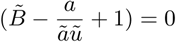, then *h*(*s*\*) is a constant function. Thus, if *h*(0) < *g*(0), then *h* and *g* have no positive intersection point. Otherwise *h*(0) > *g*(0), then h and g have at most one positive intersection point, 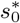. Yet, to guarantee the existence of a positive equilibrium, we must have 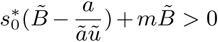. Notice that 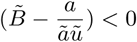.

In Fig.C.1, the graphs *h*(*s*\*) and *g*(*s*\*) intersect at two positive value of *s*\* (displayed by red dots). In the smaller inserted figure, the corresponding upper equilibrium (ordered by the size of *s*\*), 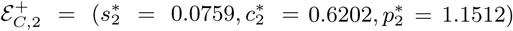, is unstable and the lower equilibrium, 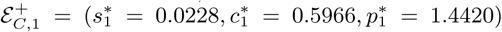, is locally asymptotically stable.

By the equality of the functions *h*(*s*\*) and *g*(*s*\*), we can also obtain *explicit solutions* for susceptible equilibria *s*\* (so can do for 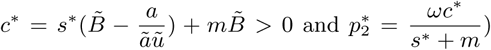.

Solving (C9) for *s*\*, we obtain

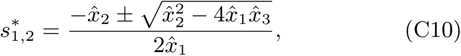

where

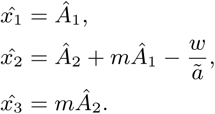

with

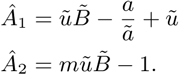

In the original non-parametrized system (III), the equation (C9) is as follows:

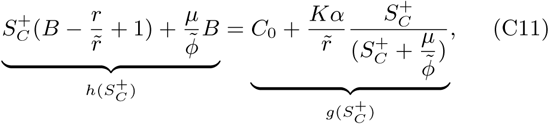

where 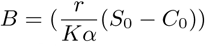.

Then we obtain

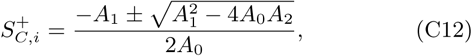

where

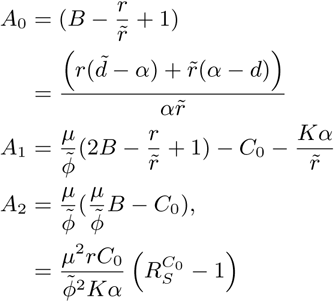

In the following subsection, we obtain the condition for backward bifurcation, proving that with vertical transmission 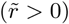, the disease outcomes change significantly.

### 3. Bistable Dynamics

Previously, we derive local stability conditions for the infection-free equilibrium, 𝓔_0_, and chronic-only equilibrium, 𝓔_c_ = (0, *C_0_, V_c_*) and point out that when both local stability conditions hold: 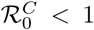 and 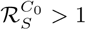, we obtain a bistable region with stable chronic-only equilibrium, ***𝓔***_*c*_ and infection-free equilibrium, 𝓔_0_. Yet, bistability can also occur with a stable interior equilibrium, 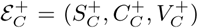, and stable infection-free equilibrium, 𝓔_0_ along with an unstable interior equilibrium or unstable chronic-only equilibrium, 𝓔_c_.

In this section, we derive a general bistability condition at the critical point 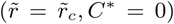, with 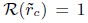, which guarantees existence of a positive interior equilibrium, 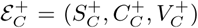 as 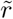 increases to 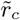. Note that

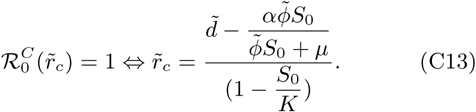

#### Theorem C.3

The chronic subsystem (III) has backward bifurcation at 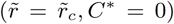 if and only if the following condition holds:

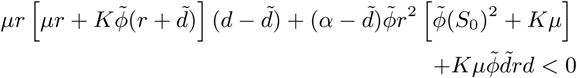

*where* 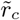 *is a critical value of the bifurcation parameter 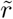 such that* 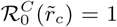. *Otherwise, if the left hand side of the condition is greater than zero, then system present forward bifurcation*.

If the chronic subsystem (III) presents a backward bifurcation, then the system dynamics are as follows:

i) the bistability

— either occurs with stable 𝓔_0_, and stable 𝓔_c_, (along with an unstable unique interior equilibrium 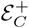),

**FIG. C.1:**
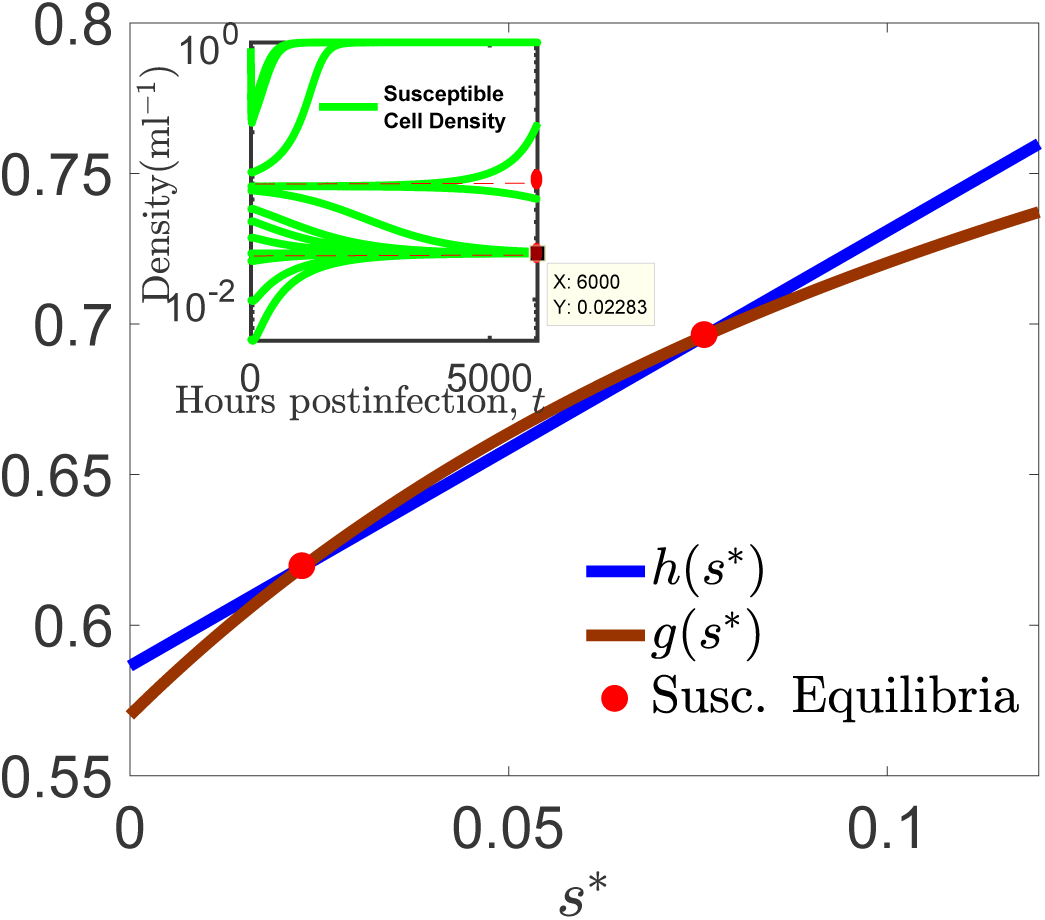
*Existence of two positive interior equilibria.* The intersection of the equations *h(s*\*) and *g(s*\*), given in (C9), provides the susceptible equilibria of the scaled system (IV). The figure above displays the case, where the system (IV) has two positive equilibria 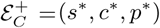 in positive orthant. Here the intersection of the two graphs, namely *h(s*\*) and *g(s*),* in larger picture pointed with red dots. In the figure, the intersections points are 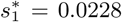 and 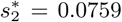 with corresponding positive equilibria 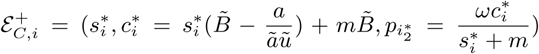. The inserted smaller figure displays time-dependent solutions of susceptible cell density with different initial conditions. The parameter values that are used here: 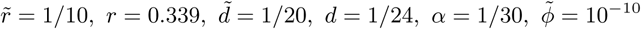, *K* = 8.947 × 10^8^, *μ* = 0.012.

— or with stable 𝓔_0_, and stable lower (or upper) interior equilibrium 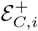 (along with unstable upper (or lower) interior equilibrium 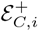),

ii) or the system only presents stable *𝓔_0_,* along with two unstable (upper and lower) interior equilibria, 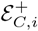 (no bistability case).

*Proof C.2 We will first show that there exists a unique continuously differentiable function 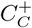 (a component of interior equilibrium* 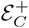), *which is a function of the bifurcation parameter* 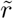 *on some open neighborhood of the critical point* 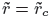. *Recall that*

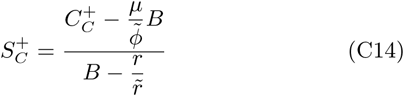

*By substituting (C14) into the equilibrium condition* 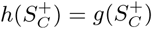, *given by (C11), we get*

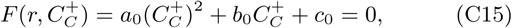

*where*

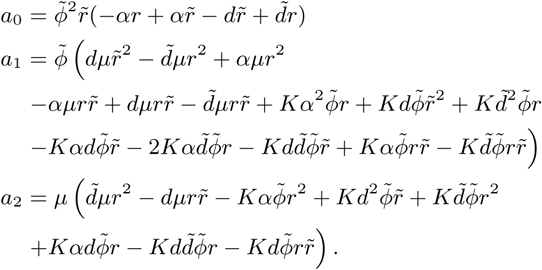

*After rearranging, we obtain, the coefficients of the polynomial (C15) as follows*:

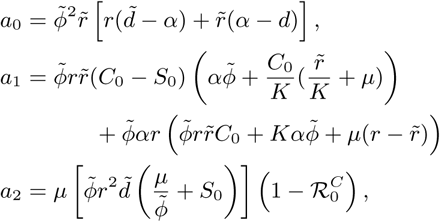

*Then at the critical point* 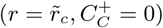, *we obtain*

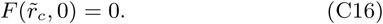

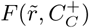 *is a continuously differentiable function of* 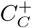. *At the fix value* 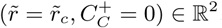, *we have*

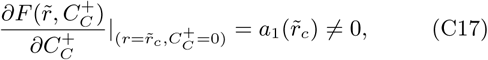

*where the expression for* 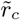 *is given by (C13) and*

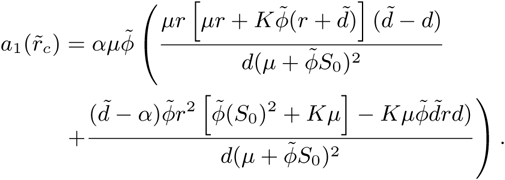

*Then by Implicit Function Theorem, there exists an open interval set 𝓤 of ℝ, containing* 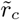, *and such that there exists a unique continuously differentiable function, f : 𝓤 →*:

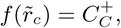

*with* 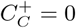, *and*

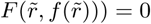

*for all* 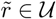. *We have an explicit expression for* 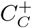 *as a function of* 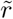, *given in (7). Then by Implicit Function Theorem, this expression of* 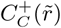 *is unique and continuously differentiable w.r.t.* 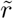 *in the open neighborhood 𝓤 of* 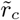.

*Next, we will investigate the sign of* 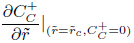. *By taking the implicit derivative of the equality (C15) w.r.t* 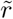 *at the point* 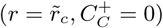, *we obtain*

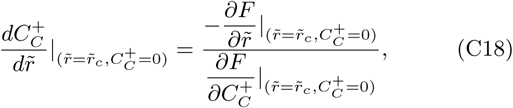

*where*

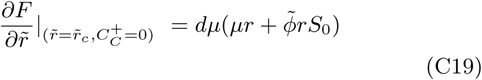

*and the expression for* 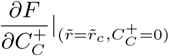 *is given in (C17)*.

*Notice that* 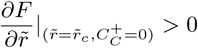. *Then*

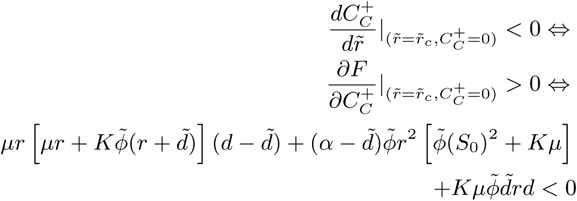

#### Theorem C.4

The chronic subsystem (III) has a backward, bifurcation at 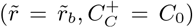 if and only if the following condition holds:

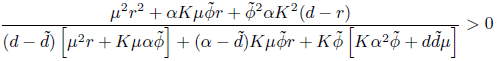

where 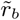 is a critical value of the bifurcation parameter 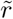 such that 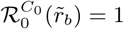. Otherwise, if the left hand side of the condition is less than zero, then the system present forward bifurcation at 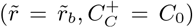.

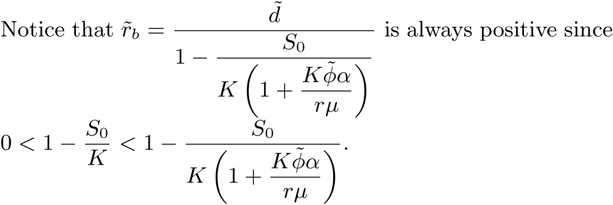

**Proof C.3** It is analogous to Proof C.2.

#### Theorem C.5

If 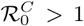, then the chronic subsystem (III) has at most a unique positive interior equilibrium 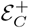. In particular, whenever

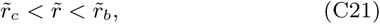

it has a unique interior equilibrium. Otherwise,

a. *if* 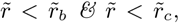, *then the system does not have a positive interior equilibrium* 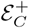, *but a stable chronic- only equilibrium, 𝓔_c_*.
b. *if* 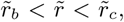, *then the system present bistability with stable chronic-only equilibrium, 𝓔_c_ and stable infection-free equilibrium*,
c. *if* 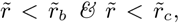, *the system has none, one or two positive interior equilibria*, 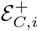 *for i = 1, 2*.

**Proof C.4** Recall the polynomials (C12), and (C15), derived from the chronic subsystem (III):

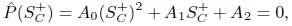

*where*

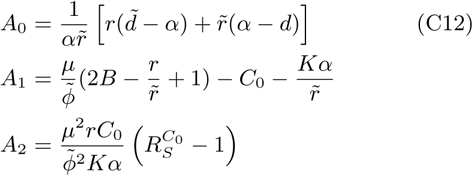

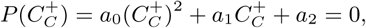

*where*

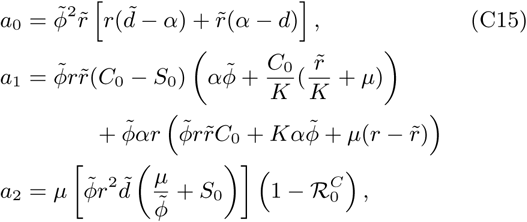

*Note that sgn[a_0_] = sgn[A_0_]:*

*Case i) Assume that* 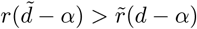.

Then the leading coefficient *a*_0_ > 0. Also note that whenever 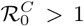, we obtain the constant term a_2_ < 0. Then, if both conditions hold: 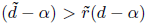, and 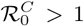, the polynomial (C15) is a concave up parabola. Then whenever 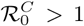, it has a unique positive root 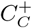, since a_2_ < 0. Therefore if 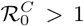, the chronic subsystem (III) has at most a unique positive interior equilibrium. Note that positivity of 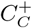 does not guarantee positivity of 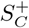 (see (C14)). Next we will establish positivity of 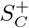 under the condition 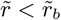.

**Claim C.1** *If* 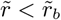, then 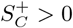.

**Proof C.5** *Note that at* 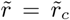, *wc have* 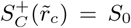. By continuity, there exists an open neighborhood, 𝓤, of 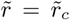 such that for all 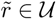, we have 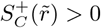. Now assume that there exists a point 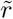 such that 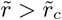 and 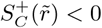. Then by intermediate value theorem, there exists a critical point 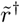: 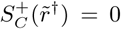. If 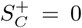, then we must have 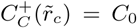. Then 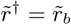, since 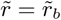 is the only critical value, providing 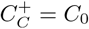.

*Case ii) Now consider that* 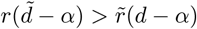.

Then the leading coefficient A_0_ < 0. Also note that whenever 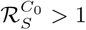, (or 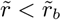) we have the constant term A_2_ > 0. Then the polynomial (C12) is a concave down parabola with constant term A_2_ > 0. Therefore if 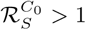, the polynomial (C12) has a positive root 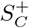.

**Claim C.2** *If* 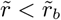, *then* 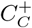 > 0.

**Proof C.6** *Since* 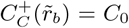, by continuity there exists an open neighborhood, 𝓥, of 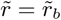: for all 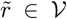, we have 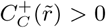. Now assume that there exists a point 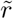 such that 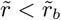 and 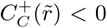. Then by intermediate value theorem, there exists a critical point 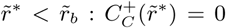. Then we must have 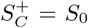. Then 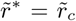 since 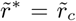 is the only critical point, providing 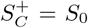.

Case iii) Now assume that

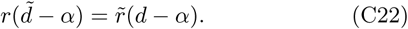

*Let* 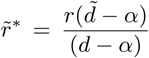 *and assume that* 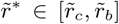. *Fix an interval* [*α_s_*, *α_e_*] *of *α* so that whenever* 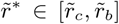, *we have α* ∈ [*α_s_*, *α_e_*]. *Let* 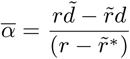. *Then whenever* 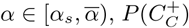 *has either one positive and one negative root, namely* 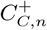 *and* 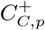, *or two positive roots, namely* 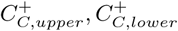. *For simplicity, assume that whenever* 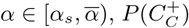 *has one positive and one negative root. Then if* 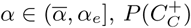 *has two poitive roots. At the point* 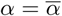, *both leading coefficients a_0_ = A_0_ = 0. By continuity, as* 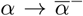, *the negative root 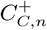 converges to -∞ and as 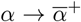, the upper positive root 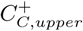 converge to +∞. Again by continuity, the continuous map 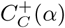, which passes through 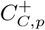 and 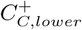, takes positive value at 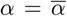; i.e. 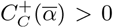. Recall that 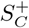 is a continuous increasing linear function of 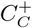 by (C14). By similar argument above, we can also show that 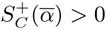*.

*Now consider that the condition (C21) does not hold. Then*

*case a. if 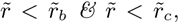, then the system does not have a positive interior equilibrium 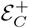, but stable chronic-only equilibrium, 𝓔_c_, since it is the parameter region of 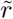 such that 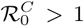 and 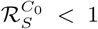*.

*b. if 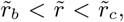, the we obtain bistability with stable chronic-only equilibrium, 𝓔_c_ and stable infection-free equilibrium, 𝓔_0_ since they are equivalent conditions to 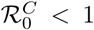, *and* 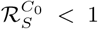*.

*case c. if* 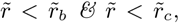, *we have sgn[A_0_] = sgn[A_2_], and, sgn[a_0_] = sgn[a_2_], for polynomials(C12), and (C15), respectively. If* 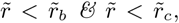, then we also have 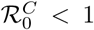, *and* 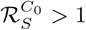. *Therefore sgn[A_0_] = sgn[A_2_] > 0. If sgn[A_0_] = sgn[A_2_] > 0,then sgn[A_0_] = sgn[A_2_] = sgn[a_0_] = sgn[a_2_]. Therefore both polynomials have either no positive roots or two positive roots. Otherwise if sgn[A_0_] = sgn[A_2_] < 0, then by Descartes’ rule of signs, it has one positive root*.

Notice that Theorem C.5 also excludes the existence of bistability or forward hysteresis (see [33]) for 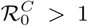.

### 4. Local Stability of the Equilibria and Hopf Bifurcation (Routh-Hurwitz Criteria)

Back to rescaled version, (IV), of the chronic system. Linearizing the system around the positive infection equilibrium 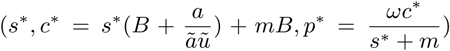, we obtain the following Jacobian matrix:

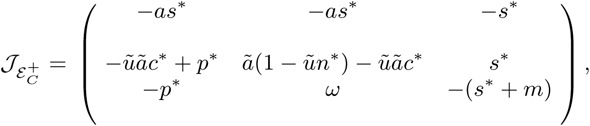

The characteristic equation for the Jacobian matrix 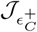 *is*:

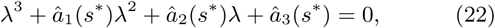

where

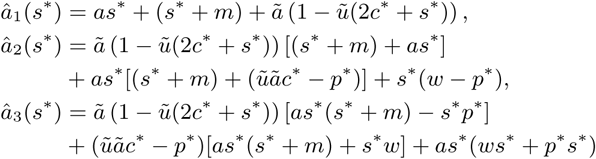

with

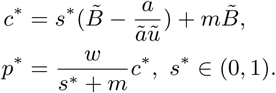

By Routh-Hurwitz Criteria, for any *s*\* ∈ (0, 1), the positive equilibrium 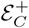 is locally asymptotically stable if and only if: 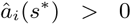, for *i* = 1, 2, 3, and 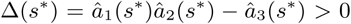. Otherwise if 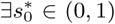 such that 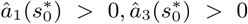 and 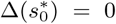, where Δ(*s*\*) is a smooth function of *s*\*, in an open interval of *s*\* : 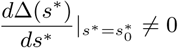, then the system exhibits Hopf bifurcation at 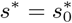.

The coefficients of the characteristic equation are functions of *s*\*, for which explicit formulas are given in (C10). Because of complicated expressions of the coefficients 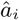, it is difficult to find any analytic condition, providing Hopf Bifurcation. Therefore we utilize numerical simulations to show that the system displays Hopf Bifurcation.

In the Fig.C.2, the x-axis presents the values of 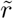 and y-axis shows how the real part of the complex eigenvalue changes w.r.t. 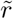. The characteristic equation, given by (22), is a cubic polynomial. So the Jacobian matrix 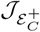 has either three real eigenvalues or one real and two complex conjugate eigenvalues, *λ*. For the case, where the Jacobian matrix 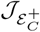 has all eigenvalues with negative real parts, the positive equilibrium 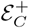 is locally asymptotically stable. If the sign of the real part of the complex eigenvalues 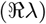 changes from negative to positive as varying the bifurcation parameter 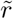, then Hopf Bifurcation occurs at 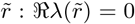. At the Hopf bifurcation point 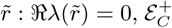 lose its stability and become unstable. Hopf bifurcation occurs when the order is reversed as well: unstable 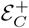 becomes stable. Notice that at 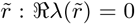, we have Δ = 0 while 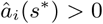, for i = 1, 2, 3. In Fig.C.2, there are two parameters values 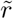 at which Hopf Bifurcation occurs. At these points, stability of equilibrium 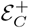 first changes from stable to unstable, and then as 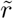 further increases, it changes unstable to stable.

## Appendix D Lytic and Chronic Invasion Analysis

Assuming that when rare lytic population arrive, the chronic type (resident) is at its equilibrium

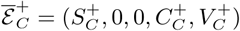

(the expressions of positive components are given in (7)), we can estimate *the lytic invasion fitness quantity 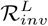* by using Next Generation Matrix Approach:

Let the entries of the matrix 𝓕 be the rates of appearance of new chronic infections among susceptible cell population in lytic infected environment, and the entries of the transition matrix 𝓥 be the rates of transfer of individuals into or out of compartments such as death, infection, or absorption.

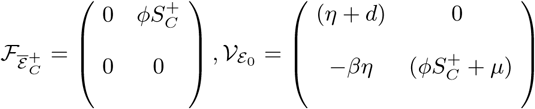

Then the lytic invasion fitness quantity is:

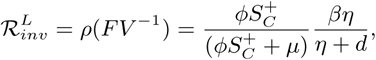

establishing the following result:

### Theorem D.1

The dominance equilibrium of chronic virus 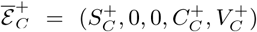 is locally asymptotically stable if 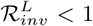 and unstable if 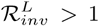.

Notice that at the chronic equilibrium 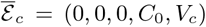 (the expressions of the positive components are given in (5)), lytic invasion does not occur due to lack of susceptible cell population in the environment and the immunity provided by the chronic infection.

By similar approach, by assuming that when rare chronic population arrive, the resident lytic population is at its equilibrium

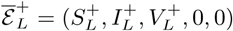

(the expressions of positive components are given in (2)), we can also estimate *the chronic virus invasion fitness quantity* as follows:

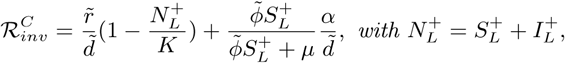

establishing the following result:

### Theorem D.2

The dominance equilibrium of lytic virus 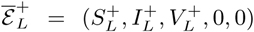 is locally asymptotically stable if 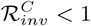 and unstable if 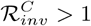.

## Appendix E Invasion Dynamics: Coexistence & Substitution

### 1. Reproduction Number of the multi-strain model

The Jacobian matrix 𝓘 evaluated at the infection-free equilibrium 𝓔_0_ = (*S*_0_, 0, 0, 0, 0) is 𝓘|*ε*_0_ = (𝓕 − 𝓥)|*ε*_0_, where

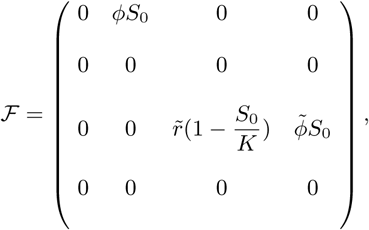

and

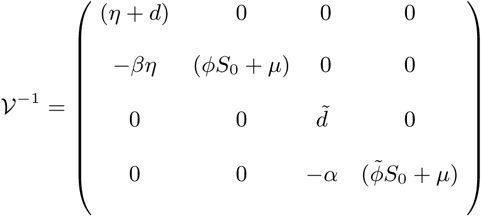

By using the Next Generation Matrix approach for the multistrain model, we obtain:

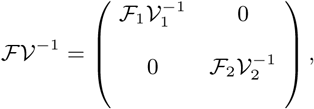

Note that 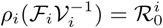. Therefore the reproduction number for the multi-strain model is

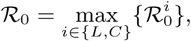

establishing the following theorem:

**FIG. C.2:**
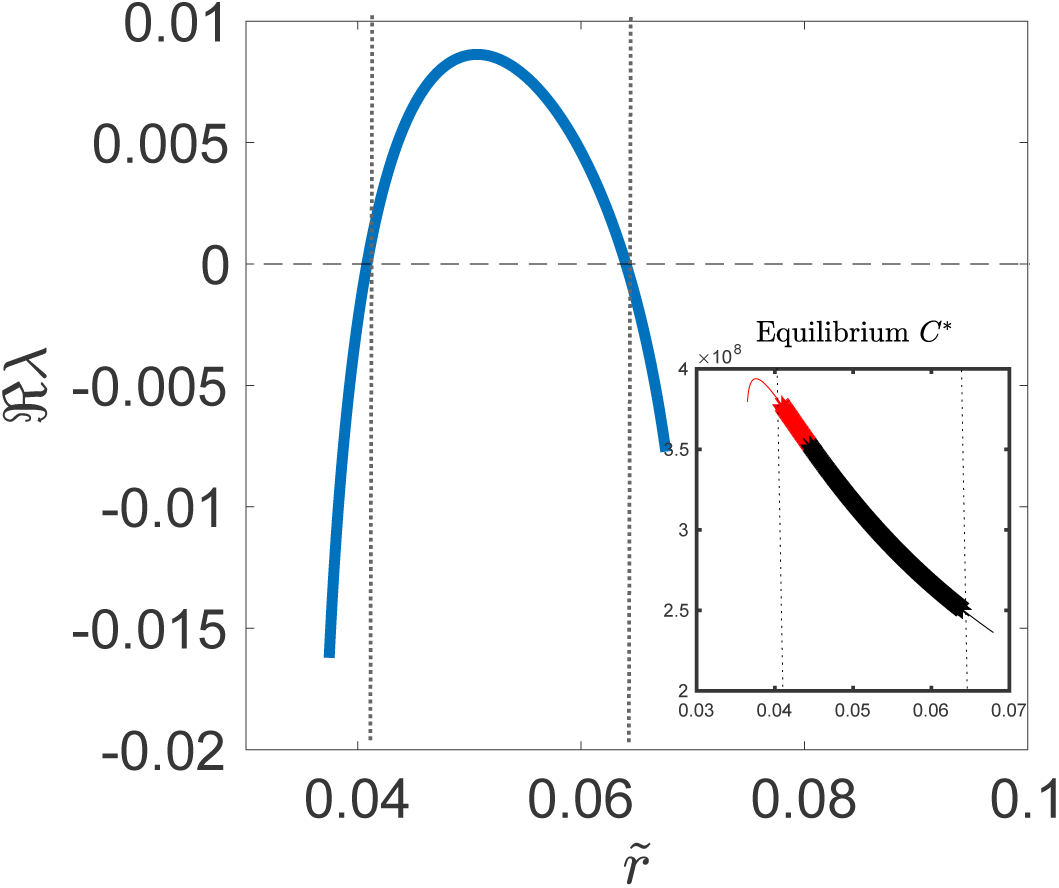
The region of Hopf bifurcation: The larger figure displays the real part of the complex eigenvalue, 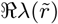 v.s. chronic virus replication rate 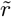. The smaller figure displays the chronic equilibrium 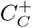 v.s. 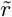. In the smaller figure, we observe that as 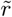 increases, locally stable positive equilibrium (displayed by red solid line) loses its stability and the system presents Hopf bifurcation, displaying sustained oscillation, shown with ★ The red color equilibrium is the upper positive equilibrium in bistabile region, observed in Fig.3(b). As 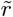 increases, the lower interior equilibrium disappear and the upper interior equilibrium become the unique positive equilibrium, displayed with black color. We obtain second Hopf bifurcation point as 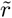 further increases. The full bifurcation diagram can be seen in Fig.3b, having the same parameter values used here.

#### Theorem E.1

If 𝓡_0_ < 1, then infection-free equilibrium *𝓔_o_* = (*S*_o_, 0, 0, 0, 0); *is locally asymptotically. Otherwise if* 𝓡_0_ > 1, *then* 𝓔_0_ is unstable.

### 2. Derivation of coexistence equilibrium for multi-strain model (I) as follows

We can rearrange the multi-strain model (I) as follows:

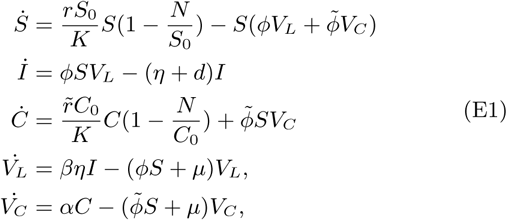

where *N* = *S* +*I* + *C*. An equilibrium, 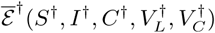, of the system (I) must be a solution of the system below:

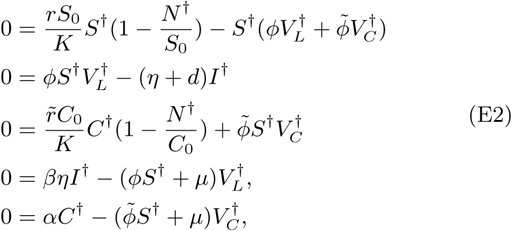

By the second and fourth equations in the system (E2), we 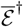, *then 𝓔_o_* have

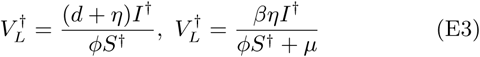

By the equality of both equations in (E3), we obtain

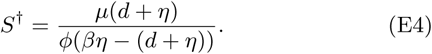

By the equation fifth equation in (E2), we have

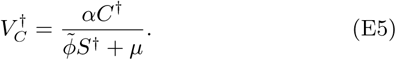

Substituting (E5), into the third equation in (E2), we get

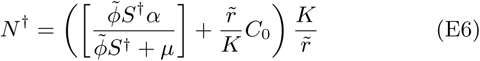

By the first equation in (E2),

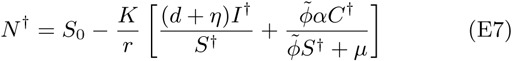

Substituting the equations (E3) and (E5) into the equation (E8), we obtain

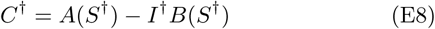

where

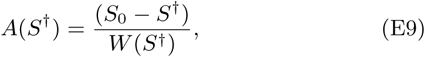

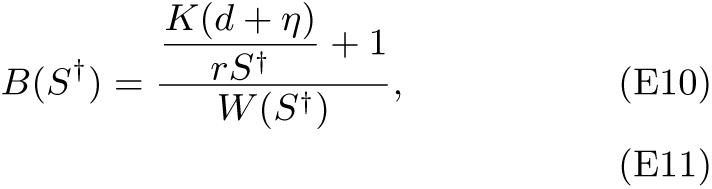

with

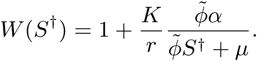

Thus

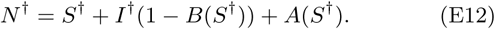

Substituting (E6) into the equation (E12) and by rearranging it, we obtain

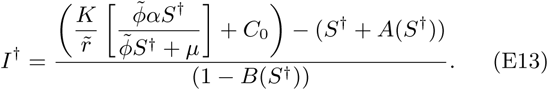

This establishes the following result:

#### Theorem E.2

The multi-strain system (I) has at most a unique coexistence equilibrium

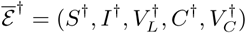

where

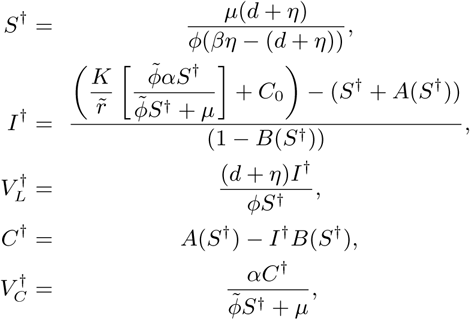

where

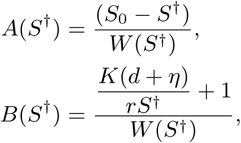

*with*

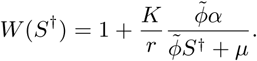

Remark E.1 Similar to the system (III), we also study the local stability of the coexistence equilibrium 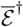 for the multistrain system (I). Evaluating the Jacobian matrix around the coexistence equilibrium 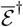, we obtain the characteristic equation, which is a fifth degree polynomial of eigenvalue λ. The coefficients of characteristic equation can be written as functions of S^†^; yet due to difficult expressions of these functions, we study the local stability of 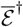, and the parameter regime, where the system exhibit hopf bifurcation, numerically. The Fig.5(d) depicts that, in the given parameter regime, the coexistence equilibrium 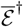 is locally asymptotically stable (displayed with •) for smaller value of *η*. Yet as *η* increases, at a critical value *η*_c_, the system undergoes hopf bifurcation and, displays sustained oscillations (the magnitude of the periodic solutions shown by a bar). The further increase in *η* restabilizes the coexistence equilibrium 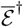.

### 3. Competitive Exclusion when 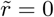

#### Theorem E.3

Assume 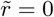. If 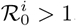, for *i* ∈ {L, C}, then the virus strain with largest reproduction number outcompetes the other one.

To prove Theorem (E.3), we will first establish the following lemma:

**Lemma E.1** *Assume* 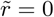. *Let*

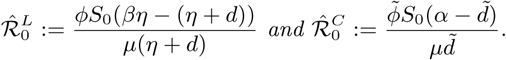

Then, for all *i* ∈ {L, C}, the reproduction numbers 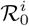 is equivalent to 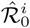 for *i* ∈ {L, C}, respectively, i.e. the following conditions hold:

i. 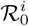 = 1 if and only if 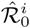 = 1,
ii. 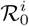 > 1 if and only if 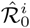 > 1,
iii. 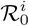 < 1 if and only if 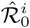 < 1.

**Proof E.1** • Case [i.] If 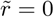, then

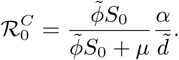

Therefore

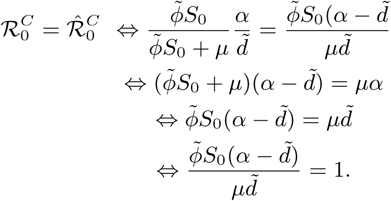

• Case [ii.]-[iii.]

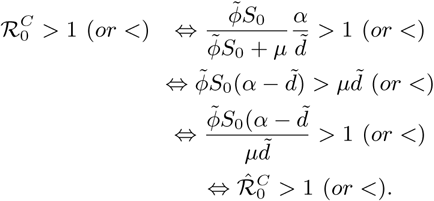

By the same argument above, one can also show that the threshold conditions 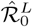 and 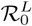 are also equivalent.

With the same argument, we can also establish the following Acknowledgments result:

**Lemma E.2** *Assume* 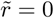. *Let*

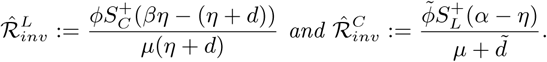

Then, for all *i* ∈ {L,C}, the invasion fitness quantity 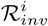 *is equivalent to* 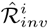, respectively.

**Proof E.2** (Proof of Theorem E.3) If 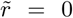, then the chronic susceptible equilibrium is

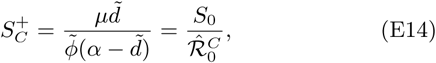

*where* 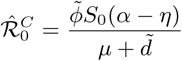. *Recall that lytic invasion conditions is*:

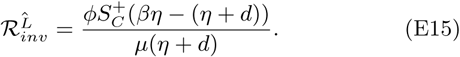

*Substituting (E14) into the equation (E15), we obtain* 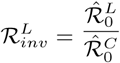. Therefore 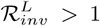 if and only if 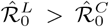 (which holds if and only if 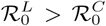.) By the same argument, we can also show 𝓡_0_ maximization for chronic virus invasion.

**TABLE V:** Infection and Invasion Regions

| Region | $\mathcal{R}_0^L$ | $\mathcal{R}_0^C$ | $\mathcal{R}_{inv}^L$ | $\mathcal{R}_{inv}^C$ | Infection | Invasion |
| --- | --- | --- | --- | --- | --- | --- |
| *I* | $< 1$ | $< 1$ | $-$ | $-$ | No infection | No invasion |
| *II* | $< 1$ | $> 1$ | $< 1$ | $-$ | Only chronic | No invasion |
| *III* | $< 1$ | $> 1$ | $> 1$ | $-$ | Only chronic | Lytic invades |
| *IV* | $> 1$ | $< 1$ | $-$ | $> 1$ | Only lytic | Chronic invades |
| *V* | $> 1$ | $< 1$ | $-$ | $< 1$ | Only lytic | No invasion |
| *VI* | $> 1$ | $> 1$ | $< 1$ | $> 1$ | Both infection | Chronic invades |
| *VII* | $> 1$ | $> 1$ | $< 1$ | $< 1$ | Both infection | No invasion |
| *VIII* | $> 1$ | $> 1$ | $> 1$ | $> 1$ | Both infection | Both invades |
| *IX* | $> 1$ | $> 1$ | $> 1$ | $< 1$ | Both infection | Lytic invades |

**FIG. E.3:**
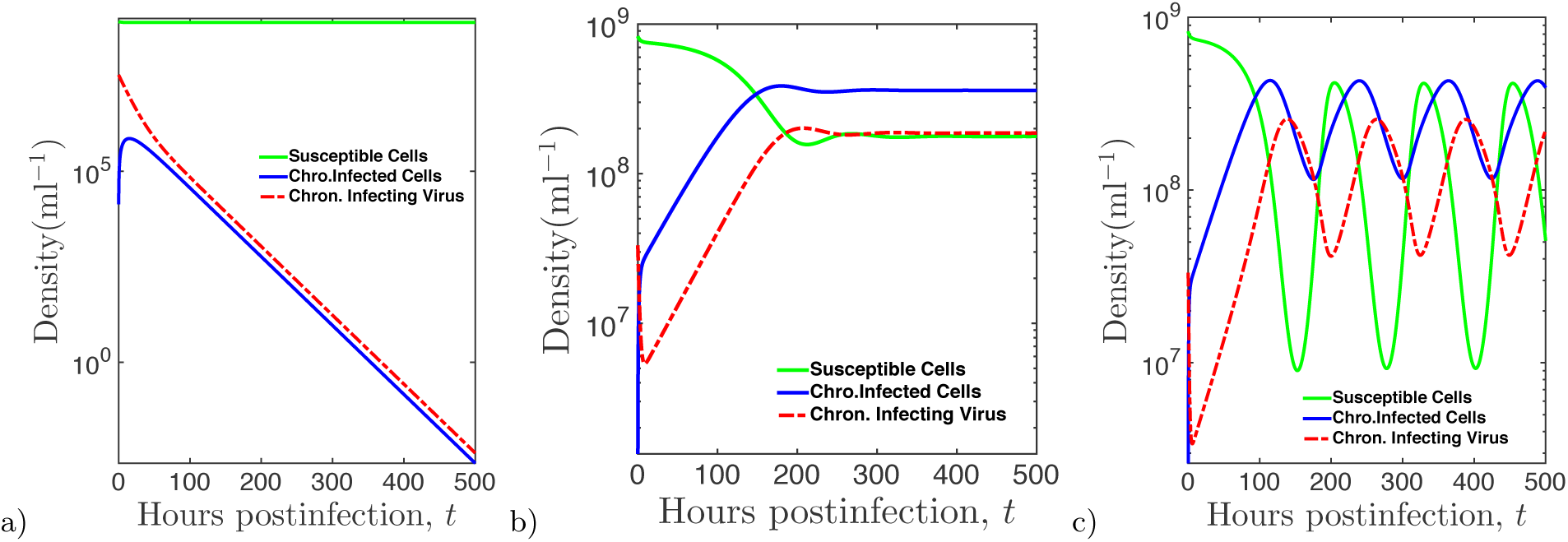
Dynamics of the chronic-subsystem (III) with susceptible, *S(t)*, chronically infected, *C(t)*, and chronically infecting free viruses, *V_C_(t)*; at time t: a) Infection dies out and population density converges to infection-free equilibrium, 𝓔_0_: b) Population density size converges to positive interior equilibrium, 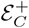. c) The system displays Hopf bifurcation: the interior equilibrium, 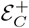 loses its stability and the system exhibits sustained oscillations. Common parameters for the dynamics are given in the Table IV. The initial virus and host densities are 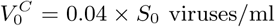 *S_0_* = 8:3 × 10^8^ hosts/ml. For part (a) 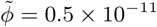, part (b) 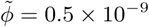 and part (c) 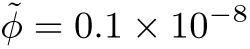.

## Acknowledgments

The authors thank Guanlin Li for helpful comments and feedback on the manuscript. JSW acknowledges support from NSF Award DEB-1342876.

---

**FIG. E.4:**
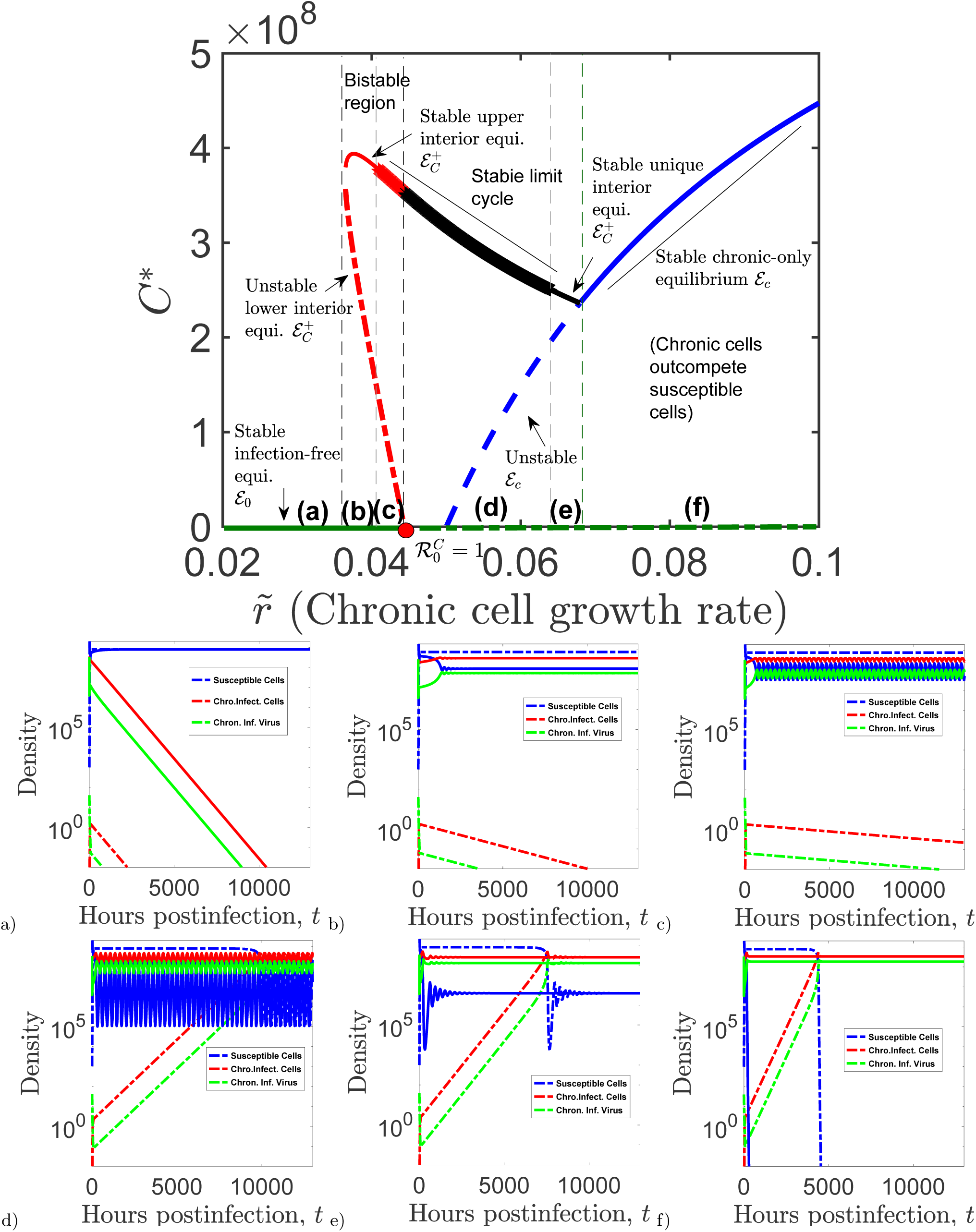
Corresponding distinct time-dependent solutions of the chronic subsystem derived from varying bifurcation parameter,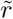. a) <u>*Region (a):*</u> Stable DFE with 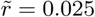) <u>*Region (b):*</u> Bistability with stable positive EE, 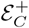, with 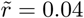. c) <u>*Region (c):*</u> Bistability with stable limit cycle with 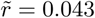) <u>*Region (d):*</u> Stable limit cycle with 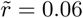) <u>*Region (e):*</u> Stable positive EE, 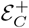, with 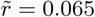) <u>*Region (f):*</u> Stable chronic-only equilibrium, 𝓔_c_, with 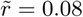. The initial virus and host densities are: *S_0_* = 8.3 x 10^9^ hosts/ml & 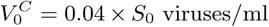 (high density); *S_0_* = 10^3^ hosts/ml, & 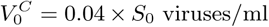 (low density), *C_0_* = 0 hosts/ml.

**FIG. E.5:**
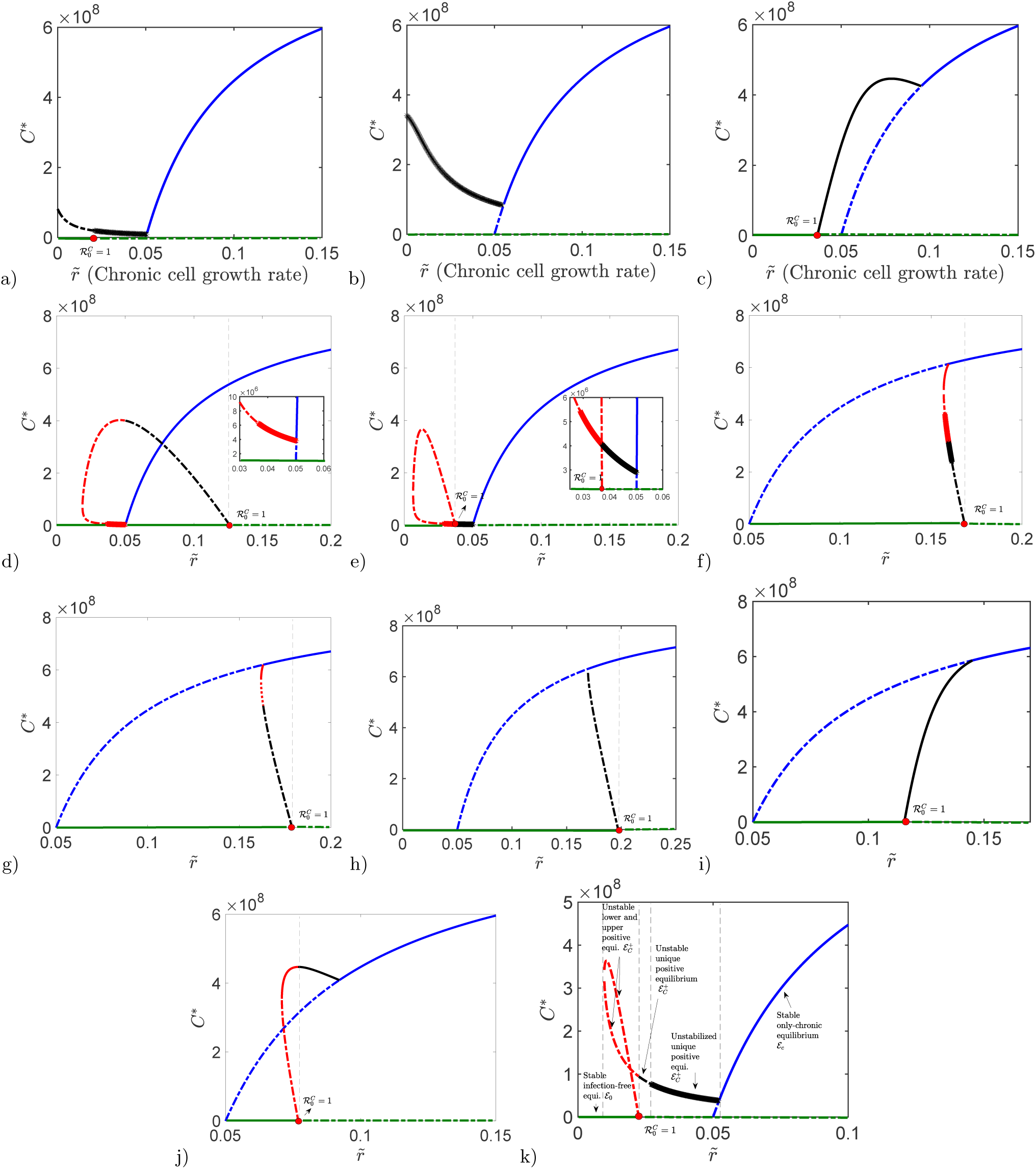
Additional bifurcation dynamics of chronic-only system (III) with varying values of the model parameters 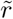 and 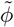. The solid (dashed) lines represent stable (unstable) equilibrium. The black lines are the unique equilibrium. The red lines shows the region, where bistability occurs (except the figure in part (k)) with stable infection-free equilibrium, 𝓔_0_ (displayed by green solid lines) and the blue lines represent chronic-only equilibrium, 𝓔_c_. The unstable infection-free equilibrium, E_0_ is displayed by green dashed lines (the region where 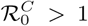). The region where the system displays sustained oscillations, shown with ★. The parameter values used here are as follows: a) *α* = 1/18, 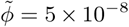, b) *α* = 1/18, 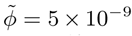, c) *α* = 1/18, 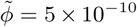, d) *α* = 1/29, 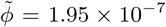, e) *α* = 1/22, 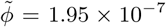, f) *α* = 1/22, 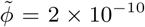, g) *α* = 1/23, 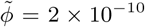 h) *α* = 1/25, 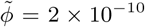, i) *α* = 1/18, 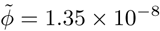), j) *α* = 1/21, φ2 = 2 x 10^−9.5^, k) *α* = 1/21, 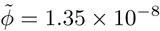, The rest of the parameter values are identical to the ones in Table I.

**FIG. E.6:**
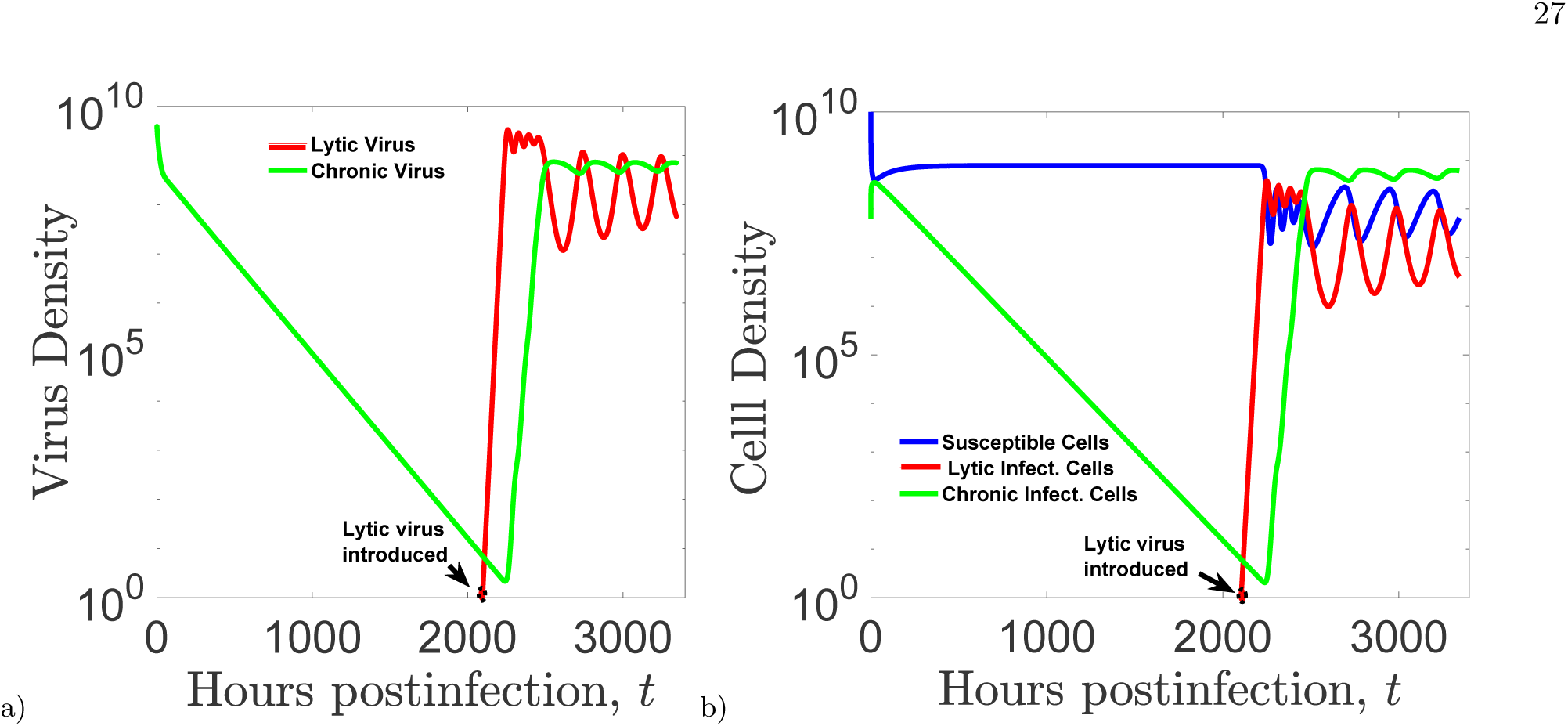
’’Rescue” of chronically infecting viruses (part a) and chronically infected cells (part b) by lytic viruses.

**TABLE IV:** Estimated parameter values of model (I)

| Parameter (Host) | Fixed Value | Source | Unit |
| --- | --- | --- | --- |
| $r$ | 0.339 | [16] | $\text{hrs}^{-1}$ |
| $K$ | $8.947 \times 10^8$ | [16] | cells/ml |
| $d$ | 1/24 | [16] | $\text{hrs}^{-1}$ |
| Parameter (Lytic) | Fixed Value | Source | Unit |
| $\phi$ | $0.88 \times 10^{-10}$ | [16] | ml/hr |
| $\beta$ | 20 | [23] | |
| $\eta$ | 0.33 | [23], [20] | |
| $\mu$ | 0.0866 | [16] | $\text{hrs}^{-1}$ |
| Parameter (Chronic) | Fixed Value | Source | Unit |
| $\tilde{r}$ | 0.2 | present study | $\text{hrs}^{-1}$ |
| $\tilde{\phi}$ | $0.2 \times 10^{-10}$ | [16] | ml/hr |
| $\alpha$ | 1/10 | present study | |
| $\tilde{d}$ | 1/20 | present study | $\text{hrs}^{-1}$ |

